# Single Cell Analysis Reveals Multiple Requirements for Zinc in the Mammalian Cell Cycle

**DOI:** 10.1101/735134

**Authors:** Maria N. Lo, Leah J. Damon, Jian Wei Tay, Amy E. Palmer

## Abstract

Despite recognition of the fundamental role of zinc (Zn^2+^) for growth and proliferation, mechanisms of how Zn^2+^ deficiency arrests these processes remain enigmatic. We induced subtle intracellular Zn^2+^ perturbations and tracked asynchronously cycling cells throughout division using fluorescent reporters, high throughput microscopy, and quantitative analysis. We found that Zn^2+^ deficiency induces quiescence and Zn^2+^ resupply stimulates cell-cycle reentry. By monitoring single cells after Zn^2+^ deprivation, we found that depending on where cells were in the cell cycle, they either went quiescent or entered the cell cycle but stalled in S phase. Stalled cells were defective in DNA synthesis and had increased DNA damage levels, suggesting a role for Zn^2+^ in maintaining genome integrity. Finally, we found that Zn^2+^ deficiency-induced quiescence does not require the cell-cycle inhibitor p21. Overall, our study provides new insights into when Zn^2+^ is required during the mammalian cell cycle and the consequences Zn^2+^ deficiency.

## Introduction

Zinc (Zn^2+^) is the second most abundant transition metal in biology and is widely recognized as an essential micronutrient to all living organisms (1). Zn^2+^ was first reported to be essential for growth of *Aspergillus niger* in 1869 and subsequently demonstrated for plants, animals, and humans (2) with the first cases of human Zn^2+^ deficiency and the associated growth and developmental disorders described in 1961 (3). Zn^2+^ deficiency has since been recognized as a global health problem, and the World Health Organization (WHO) estimates a staggering one third of the world’s population does not consume adequate Zn^2+^ and is therefore at risk for associated side effects and comorbidities (https://www.who.int/whr/2002/chapter4/en/index3.html) (4). While the clinical manifestations of Zn^2+^ deficiency are diverse and can be organism specific, one defining feature is universal: Zn^2+^ deficient cells fail to divide and proliferate normally, leading to organismal growth impairment (5). Despite recognition of the fundamental role of Zn^2+^ for proliferation, the mechanisms of how Zn^2+^ deficiency leads to cell-cycle arrest at the cellular and molecular level remain poorly defined.

Eukaryotic cell proliferation is governed by the cell-division cycle, a series of highly choreographed steps that involve gap (G1), DNA replication (S-phase), gap (G2), and mitosis (M) phases. Regulated transitions between proliferative and quiescent (i.e. reversible nonproliferative) states are essential for maintaining genome integrity and tissue homeostasis, ensuring proper development, and preventing tumorigenesis. Given the essentiality of Zn^2+^ for growth and proliferation, a fundamental question is whether Zn^2+^ serves as a nutrient, like amino acids, whether it affects the rate of cell cycle progression, or whether it is required at a specific phase of the cell cycle. Pioneering work by Chesters et al sought to define precisely when Zn^2+^ is required in the mammalian cell cycle. By chelating Zn^2+^ at different timepoints after release from serum starvation-induced quiescence, they found that Zn^2+^ was important for thymidine incorporation and thus DNA synthesis, leading to the conclusion that Zn^2+^ was required for the G1 to S transition (6). Subsequent studies confirmed that treatment of mammalian cells with high concentrations of metal chelators (DTPA and EDTA) seemed to compromise DNA synthesis (7–10). However, later studies by Chesters et al suggested that after cells passed the restriction point in mid-G1 there was no further Zn^2+^ requirement for DNA synthesis in S phase, but rather Zn^2+^ was needed to transition from G2/M back into G1 (11). The restriction point is classically defined as the point at which cells commit to completing the cell cycle, despite the presence of mitogens and/or serum (12). Thus, while these early studies suggested that Zn^2+^ was important for progression of the mammalian cell cycle, the precise role of Zn^2+^ and whether it is required at a specific stage have remained enigmatic.

There are three limitations of these early studies on the role of Zn^2+^ in cell proliferation. First, because the analyses were carried out on populations of cells, the cells were synchronized by artificial means (serum starvation or hydroxyurea treatment) and the cell cycle phase was inferred based on release from the cell cycle block. Recently, it has become clear that synchronization can induce stress response pathways that are specific to the type of arrest (13, 14). Further, cells induced into quiescence by different mechanisms (serum starvation, loss of adhesion, contact inhibition) exhibit overlapping but distinct transcriptional profiles, suggesting that different synchronization approaches impact cell cycle analysis upon emergence from quiescence (15). Second, population level analyses such as immunoblotting and qPCR mask cellular heterogeneity and subpopulations of cells with different cell fates and cell cycle dynamics (14, 16). Recent application of imaging and measurement tools for single cell analysis has uncovered distinct subpopulations of cells with different cell cycle dynamics(16), and revealed key orders of molecular events in the decision between proliferation and quiescence (16–19). Third, many of the previous investigations into the role of Zn^2+^ in the mammalian cell cycle have relied on high concentrations of chelators (DTPA, EDTA or TPEN) to induce Zn^2+^ deficiency. However, these studies did not explicitly define how these perturbations changed the intracellular labile Zn^2+^ pool, nor did they characterize how the chelators affected cell viability. Indeed, the concentrations of chelators used have been shown to induce apoptosis in a number of different cell types (20–30).

In this study, we revisit the fundamental and unresolved question of how Zn^2+^ deficiency blocks cell proliferation using a combination of fluorescent reporters, high throughput microscopy, and quantitative image analysis. It is often overlooked that in addition to serving as a reservoir for mitogens, serum is also the major source of essential micronutrients including Zn^2+^, Fe^2+^, and Mn^2+^, and thus complete removal of serum also eliminates exogenous supply of these and other essential nutrients. By controlling Zn^2+^ in the media, while maintaining mitogens at levels that normally sustain proliferation, we induced subtle perturbations of labile Zn^2+^ in the cytosol from 1 pM to 210 pM and tracked asynchronously cycling cells over multiple rounds of cell division. We found that Zn^2+^ deficiency induces cellular quiescence, but not death, and Zn^2+^ resupply stimulates synchronized cell cycle reentry. By following the entry of single cells into quiescence over time after Zn^2+^ deprivation, we found that depending on where cells were in the cell cycle, they either entered quiescence after mitosis, or entered the cell cycle but stalled in S phase. Further, we determined that cells stalled in S phase were defective in DNA synthesis and had increased levels of DNA damage, consistent with previous bulk analysis studies (31–33) suggesting a critical role for Zn^2+^ in maintaining genome integrity during replication. Finally, we found that Zn^2+^deficiency-induced quiescence does not require p21, suggesting a mechanism distinct from spontaneous quiescence (16, 19), and follows a different pattern than mitogen withdrawal. Ultimately, our study provides new insights into when Zn^2+^ is required during the mammalian cell cycle and the consequences of insufficient Zn^2+^ levels.

## Results

### Nutritional Zn^2+^ levels influence cell proliferation and intracellular Zn^2+^ levels

To revisit the question of how Zn^2+^ deficiency blocks cell growth and proliferation, we leveraged tools to visualize, track and measure molecular markers using fluorescent reporters in naturally cycling cells at the single cell level. Mammalian cells are generally recognized to contain hundreds of micromolar total Zn^2+^ (34), which they are able to concentrate from the extracellular environment. The concentration of Zn^2+^ in human serum is about 12-15 μM (35) and cell culture medias typically contain 1-40 μM Zn^2+^ (36), much of which is supplied by the serum. To rigorously control Zn^2+^ availability in our growth media, we treated serum and insulin (major sources of Zn^2+^) with Chelex^®^ 100 to scavenge Zn^2+^. We then generated a minimal media (MM) containing a low percentage of serum (1.5 %) still sufficient for proliferation that contained 1 μM Zn^2+^ as determined by Inductively Coupled Plasma Mass Spectrometry (ICP-MS), which was slightly lower than the 1.8 μM Zn^2+^ found in full growth media containing 5 % serum (Supplementary Figure S1). To further manipulate Zn^2+^, MM was either supplemented with 30 μM ZnCl_2_ to generate a Zn^2+^ replete media (ZR) or 2 – 3 μM of a Zn^2+^ specific chelator, tris(2-pyridylmethyl)amine (TPA) to generate a Zn^2+^ deficient media (ZD) (37). To establish the effect of these medias on cell viability, we grew cells in the respective medias for 30 hrs and measured viability using trypan blue. We also compared TPA to N,N,N’,N’-tetrakis-(2-pyridylmethyl)ethylenediamine (TPEN), another Zn^2+^ chelator that has been widely used in studying the effect of Zn^2+^-deficiency on cell proliferation (38–40). Even at the low end of TPEN concentrations reported in the literature (3 μM), greater than 70% cell death was observed at 24 hrs, compared to 3 μM TPA with ~ 15 % cell death. When noted, an even milder ZD condition of 2 μM TPA was used and this condition resulted in only ~ 1% cell death (Supplementary Figure S2). Our results are consistent with several studies that found TPEN induces apoptosis (20–30).

To determine how defined Zn^2+^ media conditions influenced intracellular labile Zn^2+^ levels in the cytosol, we created an MCF10A cell line stably expressing a genetically encoded FRET-based sensor for Zn^2+^ (ZapCV2 (41)), grew the cells in ZD, MM, and ZR media, and measured the resting FRET ratio in individual cells. Cells grown in ZD had a significantly lower average FRET ratio than cells grown in MM, while those grown in ZR conditions had significantly higher FRET ratios (Figure 1A). The FRET ratio correlates with the amount of labile Zn^2+^ in the cytosol, and in situ calibration suggests the respective Zn^2+^ levels are approximately 1, 80, and 210 pM for ZD, MM, and ZR media, respectively (Supplementary Figure S3), indicating that exogenous nutritional Zn^2+^ levels positively influence intracellular free Zn^2+^ levels.

**Figure 1.**
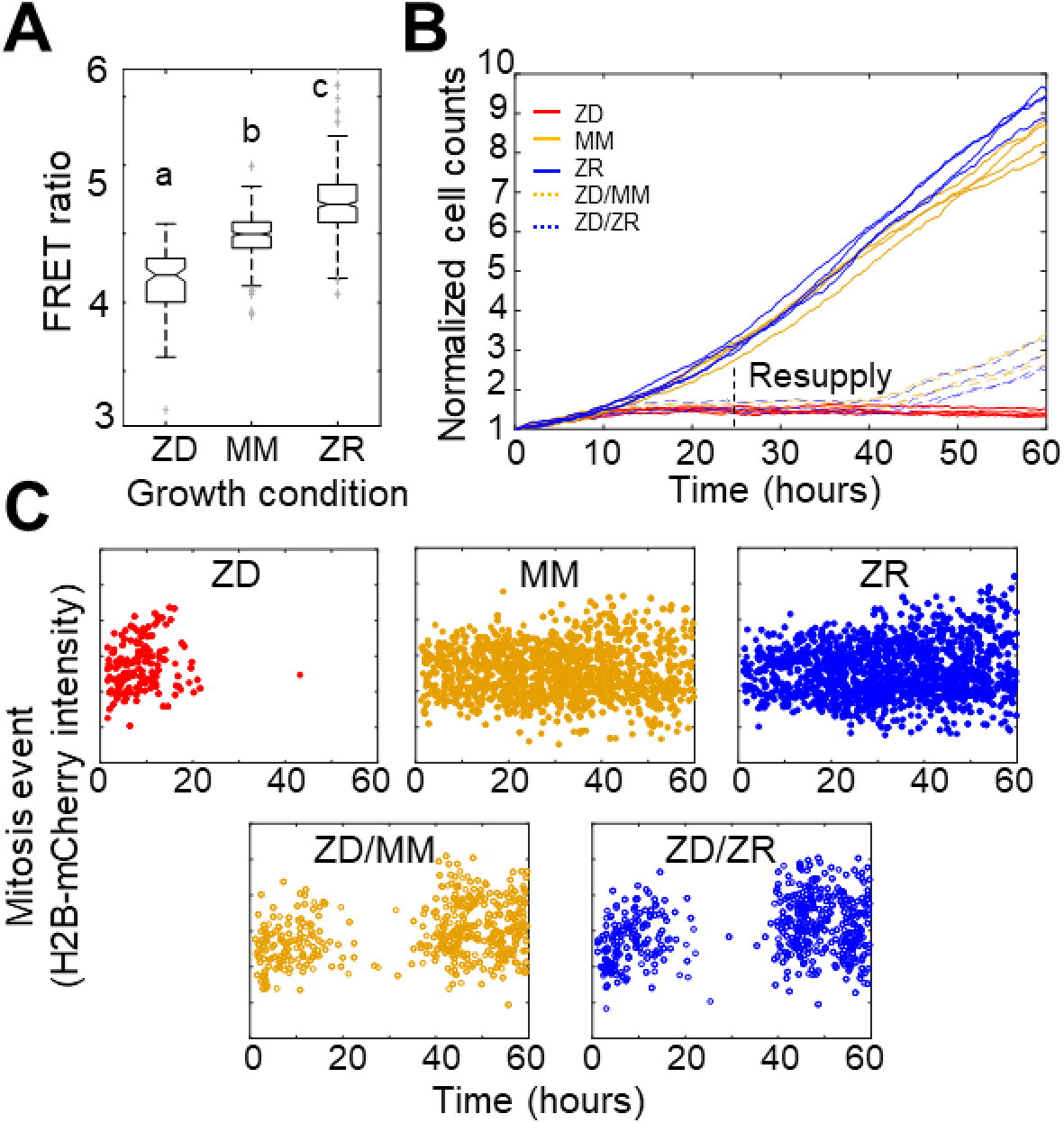
Nutritional Zn^2+^ levels influence cell proliferation and intracellular Zn^2+^ levels. A) The FRET ratio, which is proportional to labile Zn^2+^, of cells expressing ZapCV2 and grown in ZD, MM, or ZR conditions (n = 103, 382, and 402, respectively). Letters indicate statistically different groups by ANOVA with Tukey-Kramer, p < 0.001 for each comparison. DF=2 and F=333). Average FRET ratios were calculated in a one hr window before mitosis. B) Normalized cell counts after 60 hrs of growth in either Zn deficient (ZD), minimal media (MM), or Zn replete (ZR) media. ZD/MM and ZD/ZR conditions were grown in ZD media until resupply with either MM or ZR media at 24 hrs. Each line represents cells grown in an individual well (n = 4 per condition). Cell counts were normalized to initial density in each well. C) Mitosis events detected over time of cells in panel A. Mitosis events were identified as described in the Supplementary Note (Total mitosis events pooled from 4 wells).

To assess how the three nutritional Zn^2+^ regimes influenced cellular proliferation, we counted cells as a function of time. Naturally cycling cells expressing H2B-mCherry were imaged for 60 hrs and cells in each frame were segmented and counted using a custom automated analysis as described in the Supplementary Note. ZD growth conditions exhibited significantly reduced cell counts over time compared to MM and ZR conditions, with cell proliferation effectively halted after about 15 hrs (Figure 1B). Cells grown in ZR conditions reached higher cell counts, demonstrating that increased Zn^2+^ in the media promotes cellular proliferation.

Having established that our Zn^2+^ deficient conditions did not result in increased cell death, the decreased cell counts in ZD conditions could result either from a longer time between cell divisions, or an increased fraction of cells that enter a non-proliferative quiescent state. To differentiate these possibilities, we tracked individual cells over time and counted the number of mitosis events and the time between these events. Mitosis events were identified by a combination of the change in intensity of H2B-mCherry and change in size of the nucleus as described in the Supplementary Note. The number of mitosis events in ZD media decreased over time with few mitosis events detected after about 15 hrs (Figure 1C). Cells grown in MM and ZR conditions underwent mitosis events throughout the observation period, with a comparable inter-mitotic time (peaking around 13 hrs, Supplementary Figure S4). The inter-mitotic time could not be measured in ZD media because few cells underwent multiple rounds of cell division. Resupply of Zn^2+^ by adding back either MM or ZR media after 24 hrs in ZD conditions restored cell proliferation (Figure 1B, 1C), revealing that the cells were cell-cycle competent. Combined, these results suggest that mild Zn^2+^ deprivation reduces cell proliferation, not by induction of cell death, but by inducing cell cycle arrest.

### Mild Zn^2+^ deficiency induces cellular quiescence and stalling of the cell cycle at an intermediate CDK2 activity

To further examine how ZD conditions halted cell division and characterize the state of cells in ZD media, we examined single cell fate using a fluorescent reporter of CDK2 activity (16). Following cell division, CDK2 activity is low and the fluorescent reporter is localized in the nucleus, but as the cell cycle proceeds CDK2 activity increases and the reporter is progressively translocated into the cytosol (16). Thus, the ratio of cytosolic/nuclear fluorescence can be used as a readout for CDK2 activity and serves a ‘molecular timer’ for progression through the cell cycle. As described previously, CDK2 activity defines subpopulations of cells with different cell fates in a population of naturally cycling cells (16, 17, 19, 42, 43). When the CDK2 ratio remains low after mitosis (CDK2^low^), a cell is classified as quiescent whereas when CDK2 activity increases above a defined threshold within 4 hrs after mitosis (CDK2^inc^), a cell is born committed to cell cycle entry (Figure 2A). A third classification has been observed, in which a cell is born with low CDK2 activity (low CDK2 ratio) but eventually ramps up activity and commits to the cell cycle (CDK2^emerge^).

**Figure 2.**
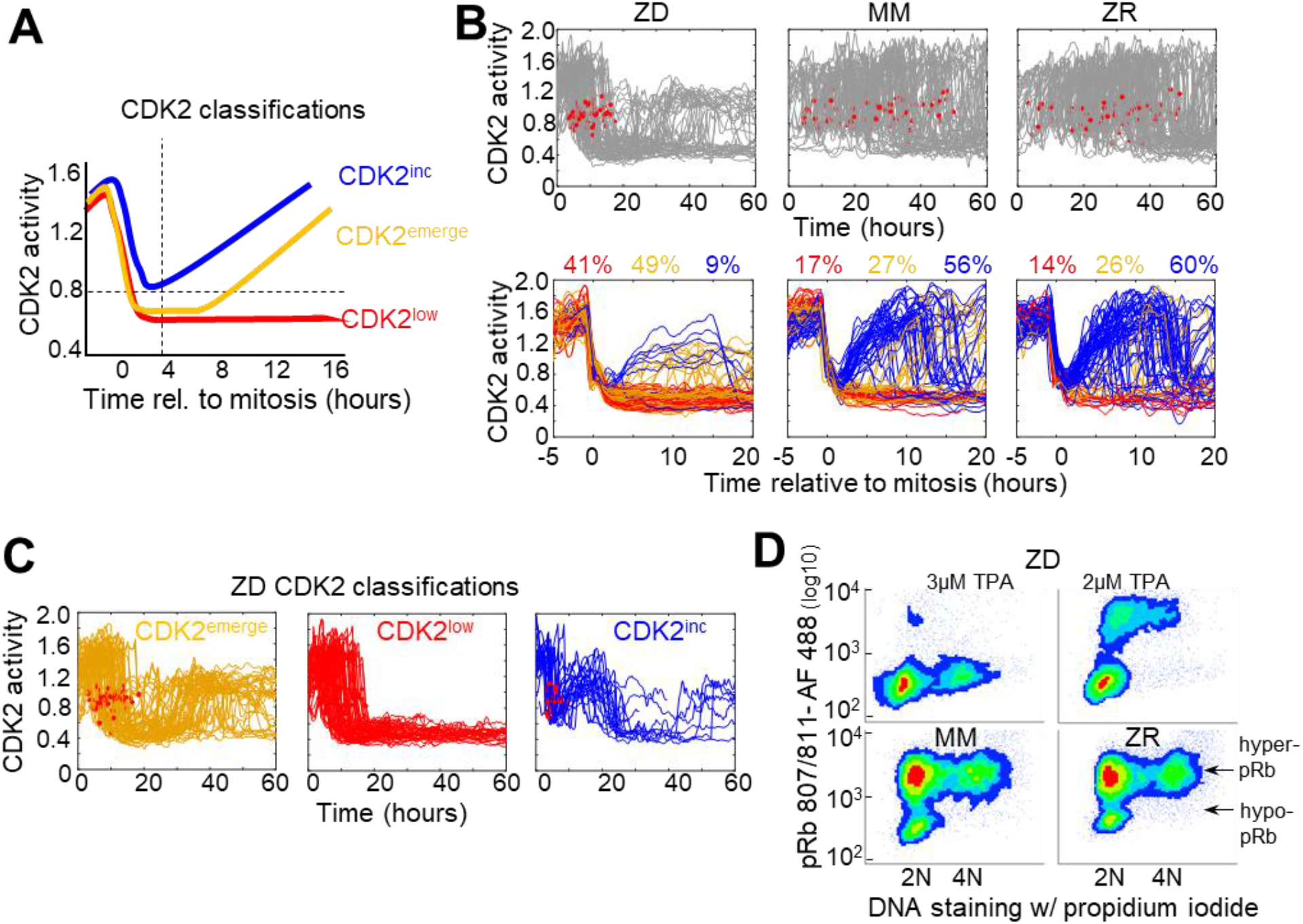
Mild Zn^2+^ deficiency induces cellular quiescence and stalling of the cell cycle in S-phase. A) Schematic of possible cell fates identified using the CDK2 activity reporter. Dashed lines indicate time and CDK2 activity thresholds used for cell fate classifications. Cell traces were computationally aligned to mitosis. B) Single cell CDK2 traces of cells grown for 60 hrs in either ZD, MM, or ZR media (n = 80 random cell traces from 4 individual wells). Gray CDK2 traces in top panel are displayed in real time and red dots indicate mitosis events. CDK2 traces in bottom panel are aligned computationally to mitosis and colored by cell fate classifications based on CDK2 activity after the first mitosis event. Percent of the total traces classified into each category are listed above traces. C) Traces from individual cell fate classifications of cells grown in ZD media as in B (n = 80 random traces). D) FACS analysis of cells grown for 24 hrs in either ZD (3 μM or 2 μM TPA), MM, or ZR media. Density scatter plots of pRb Ser 807/811 vs. PI DNA staining are displayed for each growth condition (n= 4527, 6104, 7149, 10,010, respectively). Hyper-pRb vs. hypo-pRb populations are indicated.

To define how Zn^2+^ availability in the media affects cell cycle commitment, we used an MCF10A cell line stably expressing the fluorescent CDK2 reporter and H2B-mTurquoise2 and imaged cells in ZD, MM or ZR media for 60 hrs. Individual cells were segmented, tracked, and analyzed for CDK2 activity using a custom MATLAB pipeline (Supplementary Note). In Zn^2+^ sufficient media, cells cycled naturally throughout the observation window, as evidenced by the observation of mitosis events, inter-mitotic time, and cyclical decrease in CDK2 activity after mitosis, followed by increase marking cell cycle commitment (Figure 2B, Supplementary Figure S4). When cell traces from each condition were aligned computationally to mitosis, we observed all three cell fate classifications (CDK2^inc^, CDK2^emerge^, CDK2^low^) with a similar percentage in each category in MM and ZR media (CDK2^inc^ 56% vs. 60%, CDK2^emerge^ 49% vs 41%, CDK2^low^ 17% vs 14% for MM and ZR, respectively). However, in ZD media the number of mitosis events ceased after about 20 hrs, the CDK2 activity either stayed low or rose to an intermediate level, and cells rarely underwent multiple rounds of the cell cycle (Figure 2B). Computational alignment to mitosis and cell classification revealed that few cells were born with CDK2 activity sufficient to commit to the next cell cycle (9% CDK2^inc^), a significant decrease compared to MM and ZR, and there was a substantial increase in the percentage of cells with low CDK2 activity following mitosis CDK2^low^ (41%). One of the most striking differences between ZD and MM or ZR conditions was the fate of cells that attempted to re-enter the cell cycle (i.e. whose CDK2 activity increased) following mitosis. As shown in Figure 2C, in ZD media CDK2^emerge^ cells increased CDK2 activity after a period of transient quiescence but plateaued at an intermediate CDK2 activity (a ratio of about 1.2), did not achieve maximal CDK2 activity, and failed to divide. Similarly, CDK2^inc^ cells increased CDK2 activity to an intermediate level before they dropped to a low level (Figure 2C). Analysis of mitosis events revealed that these cells did not divide before entering the CDK2^low^ state. These results suggest a significant increase in the number of cells that go quiescent after mitosis and the emergence of a new cell fate, where cells attempt to re-enter the cell cycle but stall part-way through under conditions of Zn^2+^ deficiency.

To further define how Zn^2+^ deficiency influences cell fate and characterize the consequences of the altered CDK2 activity profile in ZD cells, we examined a downstream CDK2 substrate, retinoblastoma protein (Rb). When CDK2 levels are low, Rb binds to and inhibits E2F family transcription factors, blocking cell cycle progression (44). As CDK2 activity increases, Rb gets hyper-phosphorylated which releases the inhibition, enabling E2F to transcribe cell cycle genes. A previous study showed that cells born with elevated CDK2 activity also had hyper-phosphorylated Rb (pRb), as determined by immunofluorescence, whereas cells born with CDK2^low^ had low levels of phosphorylated Rb (16). We employed a similar protocol to measure phosphorylated Rb and DNA content by Fluorescence Activated Cell Sorting (FACS). After 24 hrs of growth in MM or ZR media, the majority of cells had hyper-pRb with either 2N, intermediate, or 4N DNA content, suggesting cells were actively cycling through G1, S, G2, and M (Figure 2D). The small fraction of cells with hypo-pRb and 2N DNA content, correspond to the small fraction of cells with CDK2^low^ and represent quiescent cells. Treatment of cells with 2 μM TPA for 24 hrs revealed that most cells had 2N DNA content and hypo-pRb, consistent with most cells being in a quiescent state. However, some cells had hyper-pRb, indicating elevated CDK2 activity and an attempt to progress through the cell cycle, although there was a decrease in the fraction of cells with 4N DNA content indicating a deficiency in DNA replication. With 3 μM TPA the majority of cells had hypo-pRb with 2N DNA content, consistent with a quiescent state. A small population of cells had hypo-pRb and 4N DNA content, suggesting that after DNA replication, the cells entered quiescence without undergoing mitosis, consistent with the CDK2^inc^ population of cells in Figure 2C that slips back to a CDK2^low^ state. Combined, these results indicate that Zn^2+^ is required for cell cycle progression and there is heterogeneity in the cellular response to Zn^2+^ deprivation; some cells are born with low CDK2 activity and immediately enter quiescence, while others are born with elevated CDK2 activity (CDK2^inc^) or increase CDK2 activity after some delay (CDK2^emerge^). Further, our results suggest that the milder the Zn^2+^ deficiency, the more cells attempt to progress through the next cell cycle following mitosis. However, there is a clear requirement for Zn^2+^ to successfully progress past S-phase to G2/M, which we explore below.

### Timing of Zn^2+^ removal with respect to the cell cycle state influences cell fate

The experiments in Figure 2 revealed heterogeneity in cell fate in response to Zn^2+^ deficiency. Given that the cells were cycling asynchronously prior to Zn^2+^ deprivation, we wondered whether the cell fate was determined by a cell’s position in the cell cycle at the time of Zn^2+^ withdrawal. To address this, we imaged cells expressing the CDK2 sensor in MM for 8 hrs to track cell cycle progression prior to Zn^2+^ deprivation and follow entry of cells into quiescence. We binned cell traces according to when cells divided within specific 4 hr windows relative to TPA addition (hr 0), from 4 hrs before TPA addition (−4 to 0) up to 16 hrs after TPA addition (Figure 3A, gray shaded boxes). For cells that divided within 4 hrs prior to TPA addition, a small but elevated percent of cells went quiescent (15% vs. 7% in MM), suggesting that cells need Zn^2+^ when exiting mitosis and progressing into G1. Still, when cells divided within 4 hrs of Zn^2+^ deprivation, the majority of cells re-enter the cell cycle either immediately (CDK2^inc^) or with a slight delay (CDK2^emerge^). What was striking about this population of cells, was that only a small fraction was able to complete the next round of cell division compared to cells in MM conditions (Figure 3A top 2 panels), and instead most cells stalled with an intermediate CDK2 ratio. Thus, even if cells are born with elevated CDK2 activity and pass the classical restriction point defined by a need for extracellular growth signals such as mitogens(12), they rarely progress past an intermediate CDK2 activity in the absence of Zn^2+^. In cells that divided in subsequent windows of time after TPA addition, there was a progressive decrease in the percentage of cells born with elevated CDK2 activity and classified as CDK2^inc^ (43%, 30%, 17%, and 5%), indicating that longer Zn^2+^ deprivation increases the probability of cells entering quiescence after mitosis (Figure 3A top to bottom). There was a small increase in the percent of quiescent cells in unperturbed MM over time due to increased cell density and quiescence induced by contact inhibition. Notably, when Zn^2+^ was removed prior to cell division and cells attempted to enter another round of the cell cycle, they stalled at an intermediate CDK2 activity, consistent with an inability to progress past S-phase in the absence of Zn^2+^.

**Figure 3.**
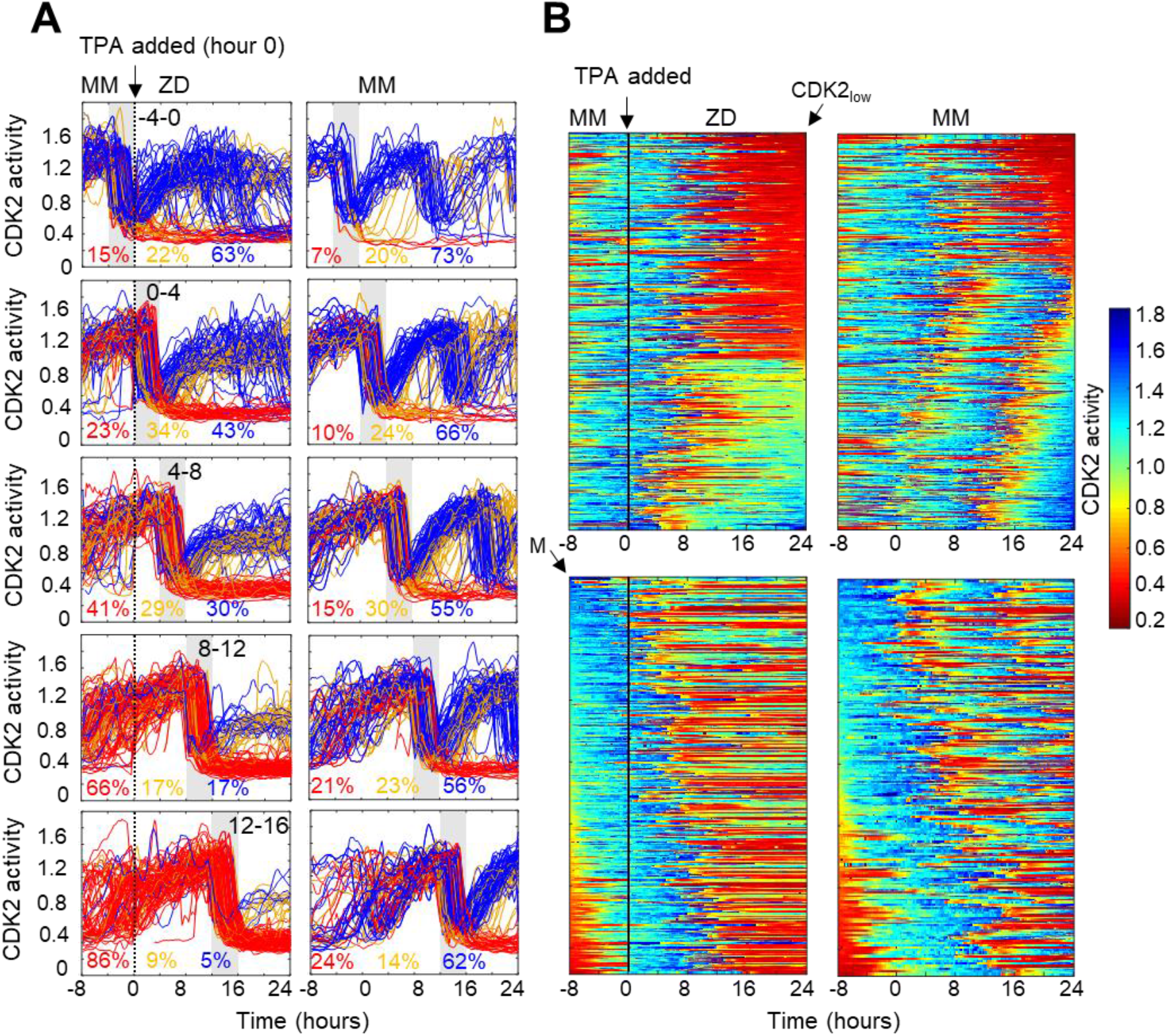
Timing of Zn^2+^ deprivation with respect to cell cycle phase influences cell fate. A) Single cell traces of CDK2 ratio over time in cells grown for 8 hrs in MM followed by 24 hrs in ZD conditions (left panels) or in unperturbed MM (right panels). Arrow/black vertical bar indicates time of TPA addition. Traces are separated by windows of when they underwent mitosis events relative to TPA addition. These 4 hr windows are shaded in gray (−4 to 0 hrs relative to TPA addition, 0-4, 4-8, 8-12, and 12-16). Each plot shows n = 120 random traces (except the top right plot: n=43 due to fewer cells identified in window) from 24 wells for TPA perturbed cells and 8 wells for MM cells. The % traces in each category are listed below each plot. Number of cells analyzed for these calculations: n=120-760 except top right window n=43). Cells are classified and color-coded based on the CDK2 ratio immediately following the first mitosis event. B) Heatmaps of CDK2 traces from individual cells as grown in A, sorted computationally. Top panel heatmaps are sorted from CDK2_low_ to CDK2_high_ at end of the imaging period. Bottom panel heatmaps are sorted by the time of mitosis (M) at the start of the imaging period. Legend shows the CDK2 color heatmap. n = 321 traces for each heatmap.

To more precisely examine the timing of events (Zn^2+^ removal, mitosis and cell fate) and compare to the behavior to mitogen removal, we plotted heat maps of individual cell traces over time (Figure 3B). We computationally grouped cell traces by their CDK2 activity at the end of the growth period, demonstrating 1) the two major cell fates for ZD cells: quiescence in red and stalled at intermediate CDK2 in green-turquoise and 2) that if cells divided more than 8 hrs after TPA addition they entered quiescence immediately after cell division (Figure 3B top panel). If cells divided within 8 hrs of TPA addition, the majority re-entered the cell cycle and stalled at an intermediate CDK2 activity (green-turquoise), consistent with an inability to progress past S-phase. As the time between TPA addition and cell division increased from 0 to ~ 8 hrs, a greater proportion of cells experienced a prolonged low CDK2 activity period (longer red streak) before re-entering the cell cycle. These results suggest that if cells experience a short window of Zn^2+^ deficiency they have a greater chance of re-entering the cell cycle; but as the length of time increases, cells experience prolonged bouts of low CDK2 activity, suggesting that Zn^2+^ plays a role in processes involved in ramping up CDK2 activity to promote cell cycle entry. In contrast, the majority of cells in MM continued to cycle throughout the measurement window.

To compare Zn^2+^ deficiency-induced quiescence to mitogen withdrawal, we aligned CKD2 traces computationally to the time of mitosis, with the first mitosis event at the top of the heat map (3B, bottom panel). Previously, this analysis demonstrated that if mitogens are withdrawn from newly born CDK2^inc^ cells, they completed one additional round of the cell cycle(16), indicating that achieving a certain threshold of CDK2 activity marks cell-cycle commitment, regardless of mitogen availability. When aligned in a similar manner (Figure 3B, bottom), our traces reveal that the CDK2 activity window that defines cell cycle commitment with respect to mitogen removal does not apply to Zn^2+^ removal. Instead, as other data representations suggest, even if cells pass the restriction point for mitogens, they stall at an intermediate CDK2 activity in the absence of sufficient Zn^2+^, suggesting that Zn^2+^ deficiency-induced quiescence acts through distinct pathways compared to mitogen withdrawal.

### Zn^2+^ deficiency causes a defect in DNA synthesis

Because we found a large population of cells which stalled at an intermediate CDK2 activity under Zn^2+^ deficient conditions and speculated that these cells were stalled in S phase, we wanted to measure whether these cells were capable of DNA synthesis. We grew cells as in Figure 3, measured 5-ethynyl-2’-deoxyuridine (EdU) incorporation during a 15 min window of labeling at the end of this growth period, and stained for DNA content using propidium iodide/RNAse. Plotting EdU intensity against DNA content for single cells revealed the expected distribution of cell cycle phases, where EdU negative cells (Edu^-^) with 2N DNA were in quiescence (G0) or G1, EdU^-^ cells with 4N DNA were in G2 or M, and EdU positive cells (Edu^+^) transitioning between 2N and 4N were in S phase (Figure 4A, cell cycle phases shown in boxes on right plot). In MM and ZR, the Edu vs. DNA content density plot followed a classical arch distribution, as has been found previously for MCF10A in full growth media (45). In ZD conditions, cells did not exhibit high EdU intensity, indicating normal DNA synthesis was impaired, and a large portion of cells were EdU^-^ with 2N DNA content, consistent with quiescence. Interestingly, in ZD conditions, many cells between 2N and 4N DNA content exhibited some Edu staining above that of cells classified as EdU^-^, suggesting that some cells are able to enter S phase, begin DNA synthesis, but at a reduced rate in the 15 min interval. The EdU intensities for cells grown in 2μM TPA were slightly higher than those for cells grown in 3μM TPA, demonstrating that 2μM TPA is a milder ZD condition and impairment of DNA synthesis was less severe. These data confirm our findings from Figure 3, where in addition to quiescence, a cell fate with intermediate CDK2 activity exists. This state of intermediate CDK2 activity is indeed S phase, as indicated by cells undergoing DNA synthesis, albeit at a reduced rate. Thus, though these cells have crossed a G1/S transition, Zn^2+^ is required in S phase for DNA synthesis to proceed at a normal rate and for cells to complete DNA synthesis and progress to G2/M.

**Figure 4.**
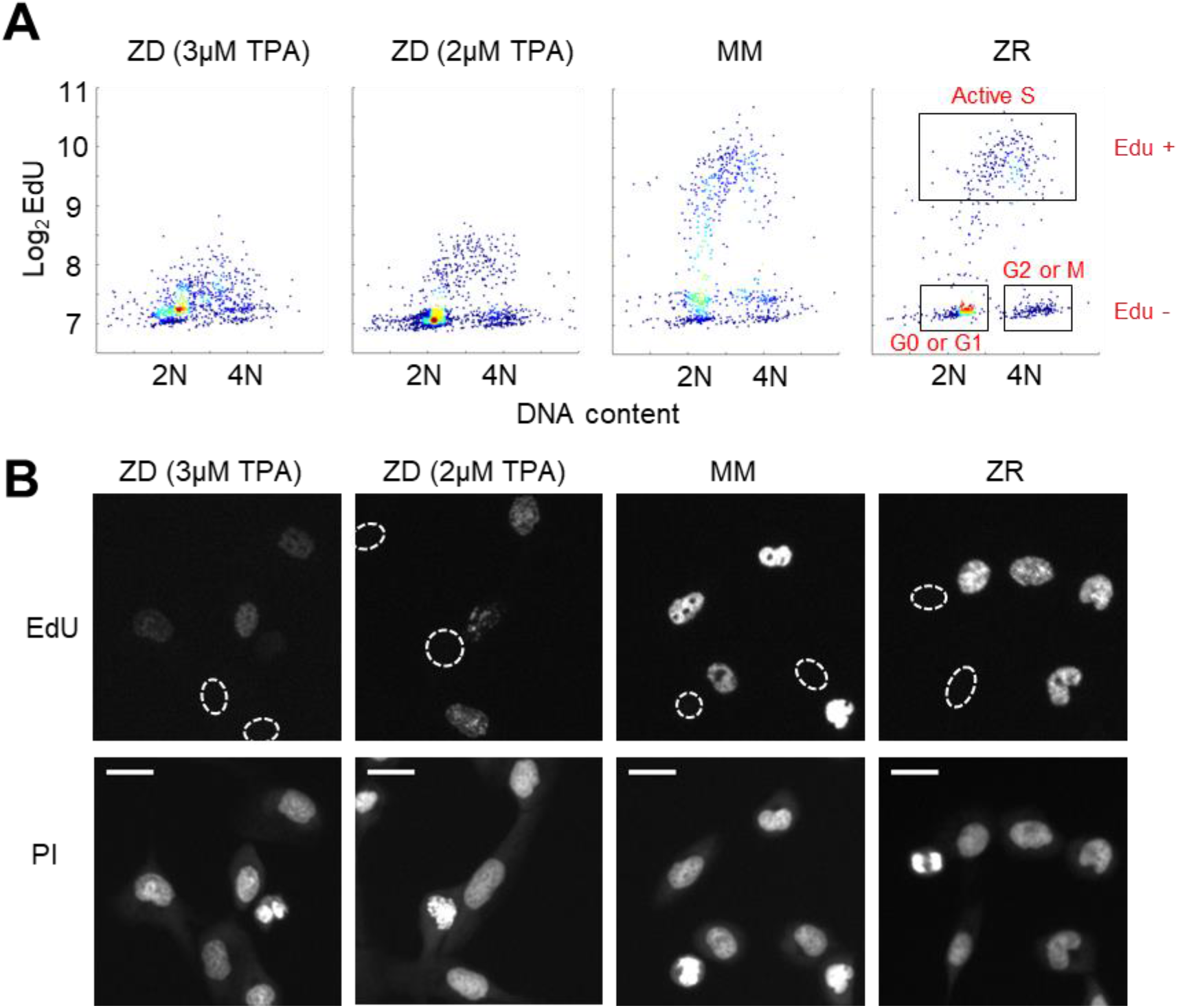
Zn^2+^ deficiency causes a defect in DNA synthesis. A) Density plots of single cells analyzed for EdU incorporation and DNA content. EdU incorporation was measured after 15 mins of incubation after 24 hrs of growth in either ZD (3μM or 2μM TPA), MM, or ZR conditions (n = 1437, 916, 817, 676, respectively from 2-4 wells per condition). Propidium iodide (PI) plus RNAse was used to stain for DNA content define 2N and 4N populations. Cell cycle phases were classified by their location on the 2D Edu vs. DNA plot (shown in red). B) Representative cell images for Edu and PI staining for each growth condition. EdU negative cells are outlined with a white dash lined. Scale bars indicate 20 μm.

### Zn^2+^ deficiency leads to increased levels of DNA damage in cycling cells

Cells that experience mild DNA damage can temporarily exit the cell cycle and enter a quiescent state in order to avoid passing a damaged genome on to the next generation. Given that long term (multi-day) growth in Zn^2+^ deficient media has previously been shown to increase DNA damage in a population of cells by the comet assay (31–33), we wondered whether mild Zn^2+^ deficiency could induce DNA damage on a shorter timescale and whether this DNA damage could be a cause of quiescence induced by Zn^2+^ deficiency. Further, because we found that ZD conditions caused a defect in DNA synthesis, we wanted to assess whether cells stalled in S phase were experiencing replication stress. Because we observed heterogeneity in cell cycle fates upon Zn^2+^ withdrawal, we sought to measure DNA damage at the single cell level and correlate it with a cell cycle marker. Previously Arora and coworkers determined that whether naturally cycling cells enter quiescence and how long they spend in quiescence is in part explained by levels of double-stranded break (DSB) DNA damage inherited from mother cells, as measured by tracking fluorescent 53BP1 foci (a known marker of DSBs) (42). Around the same time, Barr and coworkers demonstrated that both DSBs and single strand breaks (SSBs, measured by RPA2 foci) during S phase contribute to induction of quiescence (46). We used a similar approach, quantifying 53BP1 and RPA2 puncta in individual cells to identify the presence of DSB and SSB, respectively. We correlated DNA damage markers with the cell cycle using phospho-Rb status in hundreds of individual cells exposed to 24 hrs of either ZD, MM, or ZR growth conditions, where cells that were hypo-pRb were classified as quiescent, while hyper-pRb cells were classified as cycling (G1, S, G2 or M). For cells classified as quiescent (hypo-pRb), there was very little difference in 53BP1 foci (DSBs) or RPA2 foci (SSBs) between the different media conditions, although there was a slight increase in cells with DSB in ZD versus MM and ZR media (67% versus 59% and 53%, Figure 5). These results suggest that DNA damage is likely not the primary mechanism of induction of quiescence in Zn^2+^ deficient cells. However, in cycling cells, there was an increase in 53BP1 foci (84% in ZD versus ~ 50% in MM and ZR) and RPA2 foci (~ 20% in ZD versus < 5% in MM and ZR). DNA damage was measured after 24 hrs in the respective medias, and for cells in ZD, this time point corresponds to about 60% of the cells in a low CDK2 activity/quiescent state and 40% of the cells stalled at a state with intermediate CDK2 activity, consistent with S-phase (Figure 3B, 24 hr time point). Given the increase in DNA damage in cycling cells, but relatively subtle change in DNA damage in quiescent cells, we speculate that the cells with increased DNA damage correspond to the cells stalled in the cell cycle at S phase (Figure 3), and that Zn^2+^ deficiency induces a defect in DNA synthesis (Figure 4) that contributes to the inability of these cells to progress to G2/M. Our results also suggest that Zn^2+^ deficiency can induce quiescence independent of induction of DNA damage because those cells that have gone quiescent by 24 hrs do not exhibit a profound increase in DNA damage.

**Figure 5.**
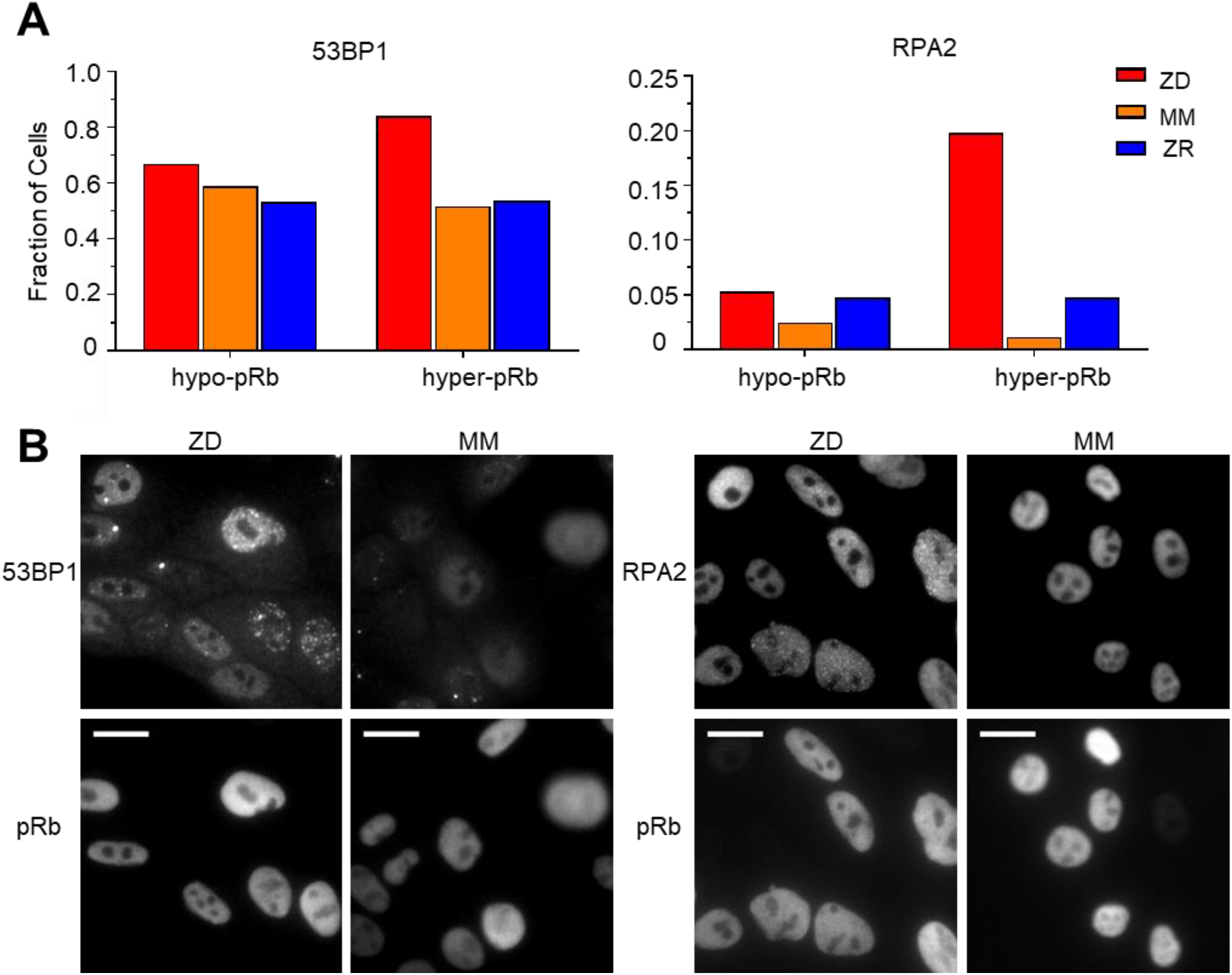
Zn^2+^ deficiency leads to increased DNA damage. A) Cells were grown in ZD, MM, or ZR media for 24 hrs and then stained with antibodies for pRB and either 53BP1 (a marker of DSB) or RPA2 (a marker of SSB). The pRB status of each cell was used to classify cells as either quiescent (hypo-pRB) or cycling (hyper-pRB). Foci were identified using a custom MATLAB script. The fraction of cells positive for DNA damage (presence of 1+ foci) is plotted. The n values for each condition are as follows: 53BP1: 1637, 1980, and 2621 for ZD, MM, and ZR; RPA2: 489, 2269, and 2439 for ZD, MM, and ZR. B) Representative images for 53BP1 and RPA2 staining for ZD and MM conditions. Scale bars indicate 20 μm.

### Quiescence induced by Zn^2+^ deficiency does not require p21

p21 is a cyclin dependent kinase inhibitor that binds to and inhibits the activity of cyclin-CDK complexes, thus regulating cell cycle progression. p21 is upregulated in response to contact inhibition and growth factor withdrawal and contributes to cell cycle arrest upon these perturbations (47). Furthermore, p21 is a transcriptional target of p53 and has been shown to be upregulated in response to DNA damage, which results in cell cycle arrest and presumably enables DNA repair prior to cell cycle re-entry. Thus, p21 has emerged as an important regulator of the proliferation-quiescence decision (19, 46). This is underscored by the observation that in p21 null cells, the incidence of spontaneous quiescence is reduced (16, 42). Given that Zn^2+^ deficiency influences cell cycle progression by inducing quiescence and that it also results in increased DNA damage, we sought to determine whether quiescence resulting from Zn^2+^ deficiency requires p21. We grew WT and p21−/− MCF10A cells expressing the CDK2 reporter and measured the fraction of cells in each classification (CDK2^low/emerge/high^) under ZD, MM, and ZR conditions (Figure 6A). In WT cells grown in MM or ZR, 17% or 16% (respectively) of cells were classified as quiescent (CDK2^low^), while in p21−/− cells, only 8% or 5% cells were classified as quiescent, consistent with previous results suggesting that spontaneous quiescence in naturally cycling cells is induced by endogenous DNA damage and is dependent on p21 (42, 46). In ZD conditions quiescence occurred at a similar rate in p21−/− and WT cells (42% vs. 56%), suggesting that quiescence caused by Zn^2+^ deficiency does not explicitly require p21 (Figure 6A). This is perhaps not surprising, given that the cell population that was quiescent after 24 hrs of growth in Zn^2+^ deficient media did not exhibit increased DNA damage (Figure 5). Combined, our data indicate that Zn^2+^-deficiency induces quiescence via a p21-independent pathway. Because p21 is also upregulated to maintain quiescence (15), we measured p21 levels in WT MCF10A cells using immunofluorescence after 40 hrs of growth. In ZD media, cells that maintained low CDK2 activity had higher levels of p21, similar to cells in MM media (Figure 6B). These data suggest that although p21 is not required for entry into quiescence, in WT cells p21 is upregulated when cells are quiescent, likely to maintain their quiescent state by suppressing CDK2 activity.

**Figure 6.**
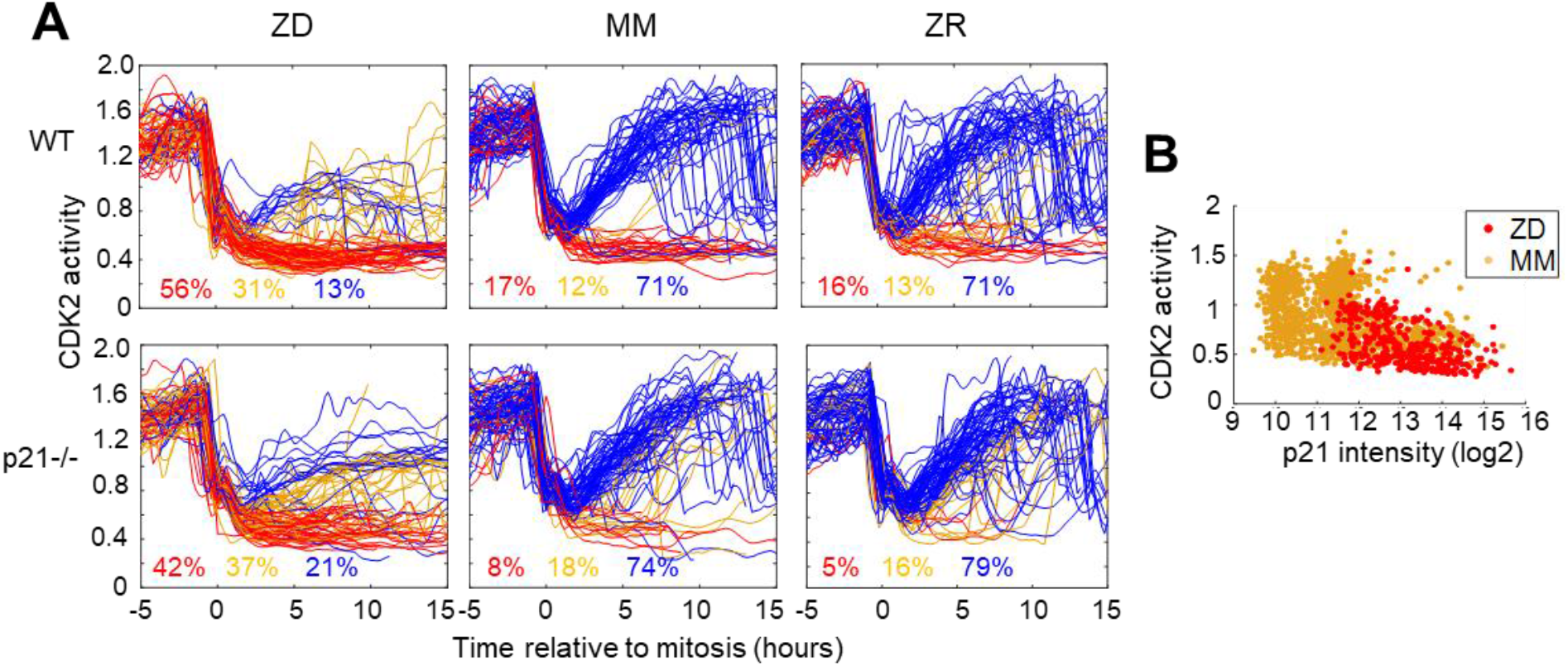
Quiescence induced by Zn^2+^deficiency does not require p21. A) Single cell CDK2 traces of WT (top) and p21−/− (bottom) MCF10A cells grown in either ZD, MM, or ZR media. Plots show n=120 random traces selected from 8 individual wells. CDK2 traces were aligned computationally to mitosis and colored by cell fate classifications. % total traces classified into each category are listed below traces (calculated from total traces n=410-3445). B) CDK2 ratio vs. p21 intensity in WT MCF10A cells at a fixed timepoint after 40 hrs of growth in ZD or MM media. p21 was detected with an anti-p21 antibody (for ZD, n = 412 cells pooled from 3 wells; for MM, n = 1414 cells pooled from 3 wells).

### Zn^2+^ resupply induces synchronized cell cycle re-entry

To determine whether Zn^2+^ deficiency-induced quiescence is reversible, we grew cells for 24 hrs in ZD media followed by 36 hrs with either MM or ZR media. Zn^2+^ resupply caused CDK2 activity to increase and the resumption of mitosis events (red dots), indicating active cell division (Figure 7A). It appeared that a smaller subset of cells remained quiescent when rescued with ZR as opposed to when rescued with MM (see CDK2^low^ traces in highlighted windows in Figure 7A). To quantify this, we generated CDK2 activity probability density plots for each hr after resupply with either MM or ZR (Figure 7B and Supplemental Figure 5). After 1 hr of Zn^2+^ resupply, the majority of cells were quiescent in all three conditions (CDK2^low^, activity mean ~ 0.5). After 8 hrs of resupply, cells not resupplied (ZD) remained in a CDK2^low^ state, while resupplied cells emerged from quiescence and entered the cell cycle, as indicated by the cell populations shifting towards higher CDK2 activity, with a mean around 1.25. The ZR resupplied cells had a higher probability of being in this higher CDK2 state and a corresponding lower probability of being in the CDK2^low^ state compared to cells resupplied with MM, suggesting a positive correlation between the amount of Zn^2+^ in the media and cell cycle re-entry after a period of deficiency.

**Figure 7.**
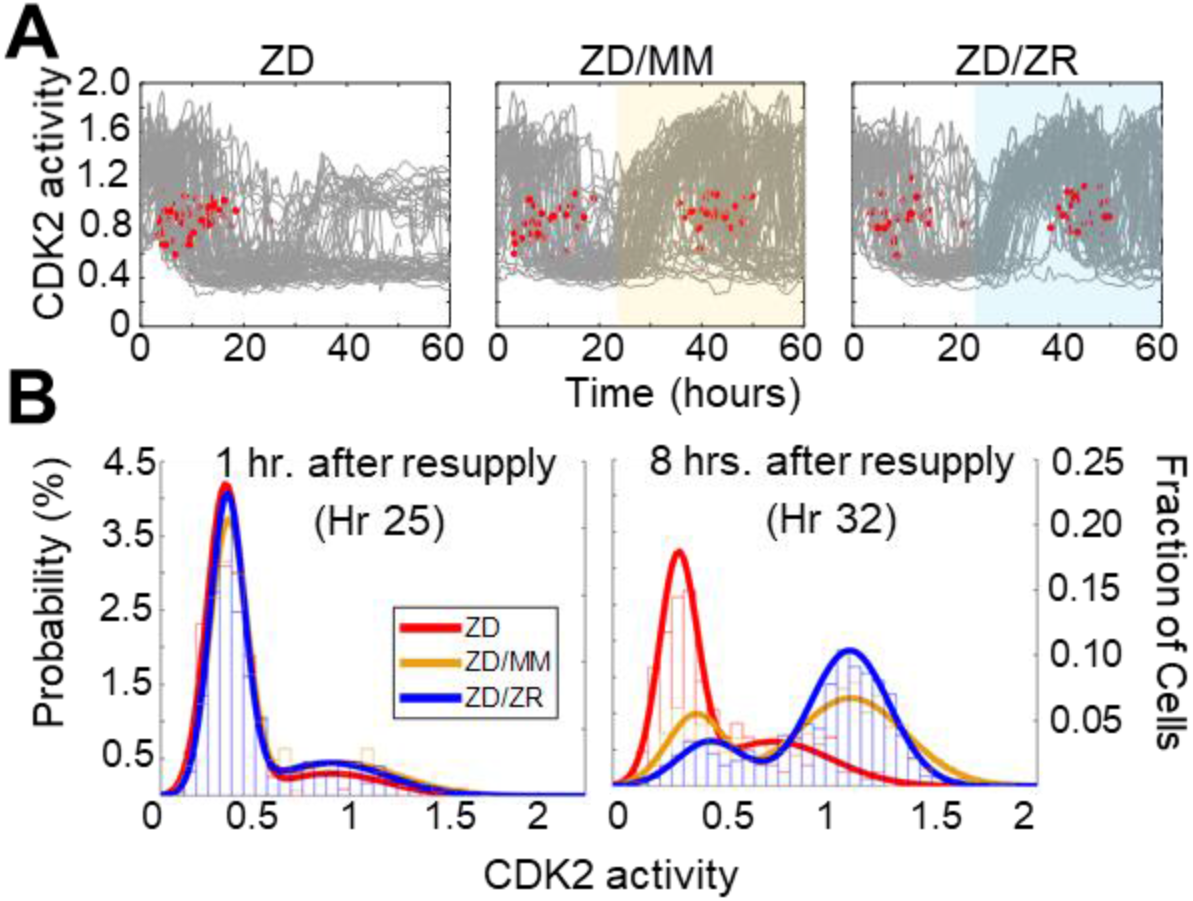
Zn^2+^ resupply after Zn^2+^-deficiency induced quiescence allows re-entry into the cell cycle. A) Single cell CDK2 traces of cells grown in ZD conditions for 60 hrs and cells grown in ZD for 24 hrs and resupplied with either MM or ZR media (n=80 random traces from 4 wells per condition). Yellow or blue shaded regions indicate media resupply. Red dots indicate mitosis events. B) CDK2 ratio density histograms after cells either remain in ZD media or are resupplied with MM or ZR. Histograms are shown for timepoints 1 and 8 hrs after resupply (n = 210, 1068, and 1024 for ZD, MM, and ZR respectively for hr 1; n = 187, 1154, and 1139 for ZD, MM, and ZR respectively for hr 8).

## Discussion

Accurate duplication of the genome and separation of chromosomes into daughter cells through the process of the cell-division cycle is one of the most essential functions individual cells must execute. Regulated exit of the cell cycle to quiescence is an important quality control pathway that reduces metabolic and biochemical activities and protects cells against stress and toxic metabolites (48). Given the importance of understanding the factors that regulate cell cycle entry, progression, and exit to quiescence, decades of research have sought to define the underlying mechanisms of the proliferation-quiescence ‘decision’. Much of our understanding of the mammalian cell cycle has derived from studies in which populations of cells were forced to exit the cell cycle upon induction of stress, such as serum starvation or amino acid deprivation. These studies have revealed many of the key regulators of the cell cycle and introduced the concept of a restriction point, a point at which cells commit to the cell cycle, and become mitogen-independent. However, recent application of single cell technologies for tracking cellular and molecular markers and the fate of individual cells have led to key revisions of the textbook model of the mammalian cell cycle, including the discovery that in naturally cycling cells there are multiple proliferation decisions (14, 16–18). Furthermore, it is now appreciated, but still poorly understood that quiescence is not a single dormant state, but rather an assemblage of heterogeneous states, that is actively maintained (14, 15, 49). Broadly speaking, control of the decision to proliferate or arrest is fundamental to various aspects of tissue architecture maintenance, differentiation, DNA damage repair, wound healing, and normal vs. cancerous cell growth (14, 18, 50). Thus, understanding how individual triggers act to induce quiescence is important for ultimately identifying targets for improving nutritional deficiencies or disease states.

Building on the emerging conceptual framework of studying the cell cycle in asynchronous populations of cells, in this work we apply a combination of fluorescent reporters, high throughput imaging, and quantitative image analysis to revisit the important but unresolved question of how Zn^2+^ deficiency blocks cell proliferation. While it has long been recognized that Zn^2+^ is required for cell proliferation, how Zn^2+^ deficiency influences the cell cycle on a single cell level has not been elucidated and whether Zn^2+^ affects the critical proliferation-quiescence cell fate decision has not been determined. Zn^2+^ is an essential metal that serves as a critical cofactor in approximately 10% of the proteins encoded by the human genome (51), including over 700 Zn^2+^-finger containing transcription factors, DNA polymerase, superoxide dismutase, and proteins involved in DNA repair (52). Thus, it is required for several key cellular processes involving these proteins, including processes relevant for the cell cycle such as transcription, antioxidant defense, DNA synthesis and repair.

It is often overlooked that in addition to serving as a reservoir for mitogens, serum is also the major source of essential micronutrients including Zn^2+^, Fe^2+^, and Mn^2+^, and thus complete removal of serum also eliminates exogenous supply of these and other essential nutrients. We sought to isolate the effect of Zn^2+^ by examining Zn^2+^ deprivation in an asynchronous population of cells still containing sufficient mitogen levels. We subjected cells to mild Zn^2+^ perturbations that altered the labile pool of cytosolic Zn^2+^ between 1 and 210 pM and avoided induction of cell death. Upon Zn^2+^ deprivation, cells lost the ability to actively proliferate and the majority experienced one of two cell fates: entry into quiescence or stall in S phase, indicating that unlike mitogens there is not a single restriction point for Zn^2+^, perhaps because it would be too hard to evolve a checkpoint for every nutrient, or perhaps because Zn^2+^ is essential for so many cellular processes. Tracking individual cells before and after Zn^2+^ deprivation revealed the temporal development of these two distinct fates, demonstrating that Zn^2+^ is required at multiple places in the cell cycle and the depth of Zn^2+^ deficiency relative to mitosis determines the cell fate. If cells underwent mitosis within 8 hrs of Zn^2+^ withdrawal, they were likely to re-enter the cell cycle and complete G1 before stalling in S phase, with a markedly prolonged S-phase (plateau at intermediate CDK2 activity in Figure 3A). These cells synthesized DNA at reduced rates and accumulated increased amounts of DNA damage, indicating that sufficient Zn^2+^ is necessary for maintaining DNA integrity. These results are consistent with previous work which showed at a population level that long term (multi-day) Zn^2+^ deficiency led to increased DNA damage as measured by a comet assay (31, 32). Here, we find that Zn^2+^ deficiency increases both DSBs and SSBs in cycling hyper-phosphorylated Rb cells (~1.7-fold increase in DSBs and ~ 4-fold increase in SSBs compared to cells in MM or ZR). It is well established that Zn^2+^ is a critical cofactor in a number of genome caretaker proteins, such as XPA and RPA of the nucleotide excision repair pathway, PARP involved in base excision repair, HERC2 involved in DSB recognition, CHFR involved in checkpoint regulation, and transcription factors such as p53 and BRCA that are involved in DNA damage response and repair (53–55). While *in vitro* studies indicate these proteins would not function without Zn^2+^, little is known about the metal-binding properties in cells, how the myriad proteins of the zinc proteome acquire their Zn^2+^, and whether these proteins are sensitive to subtle perturbations of the labile zinc pool. Ho and coworkers demonstrated that long-term Zn^2+^ depletion decreased the binding of transcription factors such as p53 and AP1 to DNA (31), and resulted in differential expression of genes involved in the cell cycle as well as DNA damage response and repair (33). However, nothing was known about how rapidly cells sense Zn^2+^ deficiency and how consequences of Zn^2+^ deficiency, such as increased DNA damage, correlate with cell fate. By following the fate of cells over time in response to Zn^2+^ withdrawal, we show that individual cells sense depletion of the labile Zn^2+^ pool quickly (within 4 hrs), and that cells in S-phase experience decreased rates of DNA synthesis and increased DNA damage, inducing cell cycle exit, despite these cells having sufficient mitogens to have passed the restriction point.

If cells divided > 8 hrs after Zn^2+^ withdrawal, the vast majority were born with low CDK2 activity and immediately entered a quiescent state. Intriguingly, these cells didn’t exhibit an increase in DNA damage and p21 was not required for entry into this quiescent state, indicating the mechanism of quiescence-induction is distinct from that of spontaneous quiescence. Entry into “spontaneous” quiescence observed in naturally cycling cells was found to be associated with increased levels of DNA damage from the previous cell cycle was dependent on p21 activity (42, 46). The observation that cells were born with low CDK2 activity suggests that Zn^2+^ deficiency is sensed in the mother cell during the previous cell cycle. This new micronutrient deficiency-induced quiescent state occurs in the presence of mitogens and is not induced via replication stress/DNA damage. Importantly, quiescent cells and cells stalled in S phase after a period of Zn^2+^ deficiency could be rescued by resupply of Zn^2+^ and were competent to re-enter the cell cycle.

By addressing the question of how mild Zn^2+^ deficiency reduces cell proliferation with modern tools at the single cell level in naturally cycling cells, we revealed that Zn^2+^ status influences multiple checkpoints in the cell cycle, and uncovered a new quiescent state induced by micronutrient deficiency that is distinct from spontaneous quiescence and mitogen withdrawal. These findings are important for understanding how this cell cycle control might be perturbed in disease states such as cancer where Zn^2+^ levels and localization have been shown to be perturbed (50, 56–58). Future proteomic and transcriptomic studies have the potential to reveal the relevant Zn^2+^ dependent proteins that sense Zn^2+^ status, thus uncovering the mechanism of entry into this quiescent state and the key Zn^2+^-dependent proteins responsible for maintaining genome integrity during DNA replication.

## Materials and Methods

### Cell culture

MCF10A cells were obtained from ATCC and maintained in full growth DMEM/F12 media (FGM) supplemented with 5% horse serum, 1% Pen/strep antibiotics, 20 ng/mL EGF, 0.5 μg/ml hydrocortisone, 100 ng/ml cholera toxin, and 10 μg/ml insulin in a humidified incubator at 37 °C and 5% CO_2_, as described previously by the Brugge lab (59). Cells were passaged with trypsin-EDTA. For imaging and growth experiments, cells were grown in 50:50 Ham’s F12 phenol red free/FluoroBrite^TM^ DMEM with 1.5% Chelex^®^ 100-treated horse serum, 1% Pen/strep antibiotics, 20 ng/mL EGF, 0.5 μg/ml hydrocortisone, 100 ng/ml cholera toxin, and 10 μg/ml Chelex^®^ 100-treated insulin. Chelex^®^ 100 was used to chelate excess Zn^2+^ from horse serum and insulin to generate the defined minimal media (MM). ZR media was generated by supplementing MM with 30 μM ZnCh and ZD media was generated by adding 2-3 μM TPA. Cell lines were routinely tested and confirmed to be mycoplasma negative by PCR.

### ICP-MS

ICP-MS was used to measure total Zn^2+^ content in defined MM with and without Chelex^®^ 100 treatment of serum and FGM. 2% nitric acid in Chelex^®^ 100 Milli-Q water was added to each media sample. Samples were spiked with 5 parts per billion (ppb) of two internal standard elements, Yttrium (Y) and Gallium (Ga) to correct for technical or human error. The ion counts of Zn^2+^ in each sample measured by ICP-MS were corrected by the ratio of measured Y or Ga (ppb) to the known amount of Y or Ga (ppb). The Zn^2+^ ion counts were then converted to ppb using a Zn^2+^ standard curve. Samples were submitted for ICP-MS analysis to the LEGS Lab at CU Boulder.

### Plasmids and cell lines

MCF10A cell lines expressing PB-NES ZapCV2 (41) and PB-H2B-mCherry were generated using the PiggyBac™ Transposon system via electroporation-mediated transfection. Stable cell lines used for long-term imaging were generated with G418 and puromycin antibiotic selection followed by FACS enrichment of dual positive fluorescent cells. The MCF10A cell line stably expressing DHB-Venus (CDK2 sensor) and H2B-mTurquoise2 and the MCF10A p21−/− were provided by the laboratory of Dr. Sabrina Spencer, CU Boulder, and generated as described previously (16). Validation of p21 knockout in this cell line was performed using immunofluorescence with a p21 antibody, as described below (See Supplemental Figure 6).

### Live cell imaging

Cells were counted with a Countess™ II Automated Cell Counter (Thermo Fisher Scientific, Waltham, MA) and plated at a density of 2,500-3,500 cells/well ~24 hrs before imaging in minimal media in glass bottom 96-well plates (P96-1.5H-N, Cellvis, Mountain View, CA). This starting density was chosen to avoid significant contact inhibition during the imaging period. In Figure 1, 2, and 5, cells were plated in 100 μL MM and 2x media of each specific nutritional regime (ZD, MM, ZR) was added immediately prior to imaging. In Figure 3, cells were plated in 100 μL MM and 2x ZD media was added after 8 hrs. In Figure 7, cells were plated in 100 μL, 2x media of specified conditions was added immediately prior to imaging; after 24 hrs, 100 μL of media was removed, and 2x media of specified conditions was resupplied. Images were collected using a Nikon Ti-E inverted microscope microscope with a Lumencor SPECTRA X light engine ^®^ (Lumencor, Beaverton, OR) and Hamamatsu Orca FLASH-4.0 V2 cMOS camera (Hamamatsu, Japan). Images were collected in time lapse series every 12 mins with a 10X 0.45 NA Plan Apo objective lens (Nikon Instruments, Melville, NY). During imaging, cells were in a controlled environmental chamber surrounding the microscope (Okolab Cage Incubator, Okolab USA INC, San Bruno, CA) at 37 °C, 5% CO_2_ and 90% humidity. Total light exposure time was ≤ 600 ms per timepoint. Filter sets used for live cell imaging and immunofluorescence (described below) were as follows: CFP Ex: 440 Em: 475/20; GFP Ex: 470, Em: 540/21; YFP Ex: 470, Em: 540/21; CFPYFP FRET Ex: 395, Em: 540/21; mCherry Ex: 555, Em: 595/40; Cy5 Ex: 640, Em: 705/22.

### Image Processing/Analysis

A detailed description of our custom MATLAB R2018A (Mathworks) pipeline for automated cell segmentation and tracking, as well as methods for calculating the FRET ratio and CDK2 ratio are provided in the Supplementary Note and the tracking code is available for download here: https://biof-git.colorado.edu/biofrontiers-imaging/palmer-zinc-cell-cycle. Briefly, mitosis events were identified when the nuclear signal generated by fluorescent H2B split into two distinct objects. The FRET ratio (FRET intensity / CFP intensity) was calculated in a cytosolic region outside the nuclear mask and the CDK2 ratio was calculated as the cytosolic intensity / nuclear intensity of the fluorescent CDK2 sensor.

### FACS

Cell pellets were washed with PBS, fixed with 4% paraformaldehyde and permeabilized with methanol at −20 °C. Cells were stained for one hr with Phospho-Rb Ser 807/811 (D20B12 XP^®^, Cell Signaling Technology, Danvers, MA) at 1:500 dilution prior to Alexa Fluor™ 488 secondary antibody (ab150073, Abcam, Cambrdige, MA) staining at 1:500 dilution for one hr. Antibodies were diluted in PBST with 1% BSA. PI DNA staining was performed using FxCycle™ PI/RNAse (Thermo Fisher Scientific F10797). FACS for phospho-Rb and PI was performed on a BD FACSCelesta™ instrument and analyzed with BD FACSDiva™ v8 software (BD Biosciences, San Jose, CA). GFP: Ex 488, Em 530/30; mCherry Ex 561, Em 610/20. FACS enrichment of stable cell lines was performed using BD FACSAria™ Fusion with the following optics: CFP: Ex 445, Em 470/15; YFP: Ex 488, Em 530/30; and mCherry Ex 561, Em 610/20.

### Immunofluorescence

Cells were fixed with 4% paraformaldehyde, washed with PBS, and permeabilized with 0.2% Triton X-100 solution. Blocking was performed in 3% BSA for one hr at 4 °C. Primary antibody staining occurred overnight at 4 °C, with the following antibody concentrations: p21, 1:200; pRB, 1:100, RPA2, 1:250; 53BP1, 1:200. Following primary staining, cells were washed with PBS and secondary antibody staining was performed for one hr with either AlexaFluor^TM^ 488, AlexaFluor™ 568, or AlexaFluor™ 647 each at a 1:500 dilution. Antibodies were diluted in PBS with 3% BSA. Hoechst staining for 15 mins at 0.1 μg/mL diluted in PBS was used to identify nuclei. For experiments comparing CDK2 activity vs. p21 intensity, the fluorescence of the CDK2 sensor was preserved upon fixation. Images were acquired on a Nikon Ti-E inverted microscope (as described in Live Cell Imaging) with a 40X 0.95 NA Plan Apo Lamda (Nikon). Primary antibodies used were: p21 Waf1/Cip1 (CST, 2947S, Lot 10), pRB Ser807/811 (CST, D20B12 XP^®^, Lot8), RPA2 (Abcam, ab2175, Lot GR3224197-5), 53BP1 (BD Biosciences, 612523, Lot 6217571) and secondaries were: AlexaFluor™ 488 (Abcam, ab150073, Lot GR328726-1), AlexaFluor™ 568 (Life Sciences A10042, Lot 1134929), or AlexaFluor™ 647 (Thermo Fisher Scientific, A-21236, Lot 1973453). Analyses were performed with custom MATLAB scripts and run in MATLAB R2018a. Briefly, nuclei were segmented using a combination of adaptive thresholding and the watershed algorithm to segment clumps of cells. The nuclear mask was used to calculate the mean nuclear pRB or p21 intensities for each cell.

For cells containing the CDK2 sensor, the nuclear mask was used to draw a ring 3 pixels wide around the nucleus for computing the CDK2 activity (defined as the ratio of cytoplasmic intensity / nuclear intensity) of the cell. For DNA damage experiments, the centroid of each cell (as defined by the nuclear mask) was used to construct a 140 x 140 pixel square around each cell; this enabled the use of MATLABs ‘adaptthresh’ function for constructing an accurate foci mask for each cell. The foci mask was further refined by filtering out any objects that were not the appropriate size (area 10-100 px^2^ for RPA2 and 10-200 px^2^ for 53BP1) or shape (eccentricity < 0.6, where 0 is a perfect circle and 1 is straight line). Cells were scored as being positive for damage when they had 1 or more focus present.

### EdU assay

Live cells kept at controlled environmental conditions (37 °C, 5 % CO_2_, 90 % humidity) were labeled with EdU according to the manufacturer’s instructions (Thermo Fisher Scientific, C10356). Briefly, cells were labeled with 10 μM EdU for 15 mins, followed by fixation and permeabilization, as described for immunofluorescence. Cells were labeled for 30 mins via a click reaction with AlexaFluor™ 647 azide using CuSO_4_ as catalyst. FxCycle™ PI/RNAse was used to quantify DNA content (2N vs 4N). Images were acquired on a Nikon Ti-E inverted microscope system with a 40X 0.95 NA Plan Apo Lambda (Nikon). Analysis was performed with custom MATLAB scripts and run in MATLAB R2018a. The analysis workflow was the same as for immunofluorescence, with the exception that high residual background EdU staining necessitated background subtraction before quantifying intensities. Background subtraction was performed by dividing the image into 11 x 11 blocks and then using the lowest 5th percentile intensity value for each block as background. Quantification of DNA content was performed by computing the integrated intensity of each cell.

### FRET sensor calibration and analysis

Sensor calibrations of MCF10A cells stably expressing PB-NES-ZapCV2 were performed using the Nikon Ti-E. Cells were grown for 24 hrs in either ZD (3 μM TPA), MM, or ZR media. For collection of R_rest_, cells were imaged for CFP-YFP FRET (200 ms exposure) and CFP (200 ms exposure) every 30 seconds for several mins. To collect R_min_, 50μM TPA in MM was added and cells were again imaged for several mins. Cells were then washed three times with phosphate, calcium, and magnesium free HEPES-buffered HBSS, pH 7.4, for removal of TPA. Finally, for collection of R_max_, cells were treated with this HBSS buffer with 119 nM buffered Zn^2+^ solution, 0.001% saponin, and 5 μM pyrithione, as previously described (60). Average R_rest_ and R_min_ were calculated by averaging across the timepoints collected. The maximum FRET ratio achieved after Zn^2+^ addition was used as R_max_. Images were background corrected by drawing a region of interest in a dark area of the image and subtracting the average fluorescence intensity of the background from the average intensity of each cell. FRET ratios for each cell (n = 8 per condition) were calculated with the following equation: (FRET_intensity_ of cell - FRET_intensity_ of background)/(CFP_intensity_ of cell - CFP_intensity_ of background). Dynamic range (DR) of the sensor in each condition was calculating as R_max_/R_min_. Fractional saturation (FS) of the sensor in each condition was calculated as follows: (R_rest_-R_min_)/(R_max_-R_min_). Finally, Zn^2+^ concentrations were estimated by [Zn^2+^] = K_D_ ((R_rest_ - R_min_)/(R_max_ - R_rest_))^1/n^, where K_D_ = 5300 pM and n = 0.29 (Hill coefficient).

### Statistical Analysis

In Figure 1A, FRET ratios between cells grown under ZD, MM, or ZR were compared using One-way ANOVA with post-hoc Tukey HSD, performed using KaleidaGraph v4.02. Alpha was 0. 05/confidence level was 0.95. Data were plotted in MATLAB v R2017b. On each box, the central mark is the median and the edges of the box are the 25th and 75th percentiles. The whiskers extend to the most extreme data points, excluding outliers, which are plotted individually with + marks.

## Acknowledgements

We thank Sabrina Spencer and members of her lab (CU Boulder) for helpful cell-cycle discussions, the CDK2 sensor cell line, and the p21−/− cell line. We thank Dr. Theresa Nahreini (Cell Culture Core Facility) for assisting with flow cytometry and Dr. Joseph Dragavon (BioFrontiers Institute Advanced Light Microscopy Core) for assisting with microscopy. This research was supported by an NIH Director’s Pioneer Award DP1-GM114863 (AEP), a Sie Foundation Postdoctoral Fellowship (MNL), and a Molecular Biophysics Training Grant T32 GM-065103 (LJD).

## Author Contributions

MNL and AEP conceptualized and designed the study. MNL and LJD performed experiments. JWT wrote cell segmentation/ tracking code and Supplementary Note. JWT, LJD, and MNL wrote image analysis and plotting scripts. MNL and LJD analyzed and interpreted data with critical feedback from AEP. MNL and AEP wrote the manuscript with edits and approval from all authors.

## Web Reference

https://www.who.int/whr/2002/chapter4/en/index3.html. Last accessed July 3, 2019

## References

1. Kaur K, Gupta R, Saraf SA, Saraf SK. Zinc: The Metal of Life. Comp Rev Food Sci F. 2014;13(4):358–76.

2. Prasad AS. Historical Aspects of Zinc. Biochemistry of Zinc Biochemistry of the Elements. 11. Boston, MA: Springer US; 1993. p. 1–15.

3. Prasad AS, Halsted JA, Nadimi M. Syndrome of iron deficiency anemia, hepatosplenomegaly, hypogonadism, dwarfism and geophagia. Am J Med. 1961;31:532–46.

4. Roohani N, Hurrell R, Kelishadi R, Schulin R. Zinc and its importance for human health: An integrative review. J Res Med Sci. 2013;18(2):144–57.

5. Vallee BL, Falchuk KH. The biochemical basis of zinc physiology. Physiol Rev. 1993;73(1):79–118.

6. Chesters JK, Petrie L, Vint H. Specificity and timing of the Zn2+ requirement for DNA synthesis by 3T3 cells. Exp Cell Res. 1989;184(2):499–508.

7. Chesters JK, Petrie L, Travis AJ. A requirement for Zn2+ for the induction of thymidine kinase but not ornithine decarboxylase in 3T3 cells stimulated from quiescence. Biochem J. 1990;272(2):525–7.

8. Chesters JK, Boyne R. Nature of the Zn2+ requirement for DNA synthesis by 3T3 cells. Exp Cell Res. 1991;192(2):631–4.

9. Watanabe K, Hasegawa K, Ohtake H, Tohyama C, Koga M. Inhibition of DNA synthesis of EDTA and its cancellation by zinc in primary cultures of adult rat hepatocytes. Biomed Res. 1993;14:99–110.

10. Prasad AS, Beck FW, Endre L, Handschu W, Kukuruga M, Kumar G. Zinc deficiency affects cell cycle and deoxythymidine kinase gene expression in HUT-78 cells. J Lab Clin Med. 1996;128(1):51–60.

11. Chesters JK, Petrie L. A possible role for cyclins in the zinc requirements during G1 and G2 phases of the cell cycle. J Nutr Biochem. 1999;10(5):279–90.

12. Pardee AB. A restriction point for control of normal animal cell proliferation. Proc Natl Acad Sci USA. 1974;71(4):1286–90.

13. Ly T, Endo A, Lamond AI. Proteomic analysis of the response to cell cycle arrests in human myeloid leukemia cells. eLife. 2015;4:e04534.

14. Matson JP, Cook JG. Cell cycle proliferation decisions: the impact of single cell analyses. FEBS J. 2017;284(3):362–75.

15. Coller HA, Sang L, Roberts JM. A New Description of Cellular Quiescence. PLoS Biol. 2006;4(3):e83.

16. Spencer Sabrina L, Cappell Steven D, Tsai F-C, Overton KW, Wang Clifford L, Meyer T. The Proliferation-Quiescence Decision Is Controlled by a Bifurcation in CDK2 Activity at Mitotic Exit. Cell. 2013;155(2):369–83.

17. Cappell SD, Chung M, Jaimovich A, Spencer SL, Meyer T. Irreversible APC(Cdh1) Inactivation Underlies the Point of No Return for Cell-Cycle Entry. Cell. 2016;166(1):167–80.

18. Heldt FS, Barr AR, Cooper S, Bakal C, Novak B. A comprehensive model for the proliferation-quiescence decision in response to endogenous DNA damage in human cells. Proc Natl Acad Sci USA. 2018;115(10):2532–7.

19. Moser J, Miller I, Carter D, Spencer SL. Control of the Restriction Point by Rb and p21. Proc Natl Acad Sci USA. 2018;115(35):E8219–E27.

20. Sunderman FW, Jr. The influence of zinc on apoptosis. Ann Clin Lab Sci. 1995;25(2):134–42.

21. Johnson VL, Ko SC, Holmstrom TH, Eriksson JE, Chow SC. Effector caspases are dispensable for the early nuclear morphological changes during chemical-induced apoptosis. J Cell Sci. 2000;113(17):2941–53.

22. Hyun HJ, Sohn JH, Ha DW, Ahn YH, Koh JY, Yoon YH. Depletion of intracellular zinc and copper with TPEN results in apoptosis of cultured human retinal pigment epithelial cells. Invest Ophthalmol Vis Sci. 2001;42(2):460–5.

23. Kolenko VM, Uzzo RG, Dulin N, Hauzman E, Bukowski R, Finke JH. Mechanism of apoptosis induced by zinc deficiency in peripheral blood T lymphocytes. Apoptosis. 2001;6(6):419–29.

24. Canzoniero LMT, Manzerra P, Sheline CT, Choi DW. Membrane-permeant chelators can attenuate Zn2+-induced cortical neuronal death. Neuropharmacology. 2003;45(3):420–8.

25. Corniola RS, Tassabehji NM, Hare J, Sharma G, Levenson CW. Zinc deficiency impairs neuronal precursor cell proliferation and induces apoptosis via p53-mediated mechanisms. Brain Res. 2008;1237:52–61.

26. Lee J-M, Kim Y-J, Ra H, Kang S-J, Han S, Koh J-Y, et al. The involvement of caspase-11 in TPEN-induced apoptosis. FEBS Lett. 2008;582(13):1871–6.

27. Makhov P, Golovine K, Uzzo RG, Rothman J, Crispen PL, Shaw T, et al. Zinc chelation induces rapid depletion of the X-linked inhibitor of apoptosis and sensitizes prostate cancer cells to TRAIL-mediated apoptosis. Cell Death Differ. 2008;15:1745.

28. Carraway RE, Dobner PR. Zinc pyrithione induces ERK-and PKC-dependent necrosis distinct from TPEN-induced apoptosis in prostate cancer cells. Biochim Biophys Acta. 2012;1823(2):544–57.

29. Mendivil-Perez M, Velez-Pardo C, Jimenez-Del-Rio M. TPEN induces apoptosis independently of zinc chelator activity in a model of acute lymphoblastic leukemia and ex vivo acute leukemia cells through oxidative stress and mitochondria caspase-3-and AIF-dependent pathways. Oxid Med Cell Longev. 2012;2012:313275-.

30. Zhu B, Wang J, Zhou F, Liu Y, Lai Y, Wang J, et al. Zinc Depletion by TPEN Induces Apoptosis in Human Acute Promyelocytic NB4 Cells. Cell Physiol Biochem. 2017;42(5):1822–36.

31. Ho E, Ames BN. Low intracellular zinc induces oxidative DNA damage, disrupts p53, NFkB, and API DNA binding, and affects DNA repair in a rat glioma cell line. Proc Natl Acad Sci USA. 2002;99(26):16770–5.

32. Ho E, Courtemanche C, Ames BN. Zinc Deficiency Induces Oxidative DNA Damage and Increases P53 Expression in Human Lung Fibroblasts. J Nutr. 2003;133(8):2543–8.

33. Yan M, Song Y, Wong CP, Hardin K, Ho E. Zinc Deficiency Alters DNA Damage Response Genes in Normal Human Prostate Epithelial Cells. J Nutr. 2008;138(4):667–73.

34. Krezel A, Maret W. Zinc-buffering capacity of a eukaryotic cell at physiological pZn. J Biol Inorg Chem. 2006;11(8):1049–62.

35. Hess SY. National Risk of Zinc Deficiency as Estimated by National Surveys. Food Nutr Bull. 2017;38(1):3–17.

36. Glassman AB, Rydzewski RS, Bennett CE. Trace metal levels in commercially prepared tissue culture media. Tissue Cell. 1980;12(4):613–7.

37. Huang Z, Zhang X-a, Bosch M, Smith SJ, Lippard SJ. Tris(2-pyridylmethyl)amine (TPA) as a membrane-permeable chelator for interception of biological mobile zinc. Metallomics. 2013;5(6):648–55.

38. Beyersmann D, Haase H. Functions of zinc in signaling, proliferation and differentiation of mammalian cells. BioMetals. 2001;14(3-4):331–41.

39. Haase H, Maret W. Intracellular zinc fluctuations modulate protein tyrosine phosphatase activity in insulin/insulin-like growth factor-1 signaling. Exp Cell Res. 2003;291(2):289–98.

40. Kaltenberg J, Plum LM, Ober-Blöbaum JL, Hönscheid A, Rink L, Haase H. Zinc signals promote IL-2-dependent proliferation of T cells. Eur J Immunol. 2010;40(5):1496–503.

41. Fiedler BL, Van Buskirk S, Carter KP, Qin Y, Carpenter MC, Palmer AE, et al. Droplet Microfluidic Flow Cytometer For Sorting On Transient Cellular Responses Of Genetically-Encoded Sensors. Anal Chem. 2017;89(1):711–9.

42. Arora M, Moser J, Phadke H, Basha AA, Spencer SL. Endogenous Replication Stress in Mother Cells Leads to Quiescence of Daughter Cells. Cell Reports. 2017;19(7):1351–64.

43. Yang HW, Chung M, Kudo T, Meyer T. Competing memories of mitogen and p53 signalling control cell-cycle entry. Nature. 2017;549:404.

44. Giacinti C, Giordano A. RB and cell cycle progression. Oncogene. 2006;25(38):5220–7.

45. Gookin S, Min M, Phadke H, Chung M, Moser J, Miller I, et al. A map of protein dynamics during cell-cycle progression and cell-cycle exit. PLoS Biol. 2017;15(9):e2003268.

46. Barr AR, Cooper S, Heldt FS, Butera F, Stoy H, Mansfeld J, et al. DNA damage during S-phase mediates the proliferation-quiescence decision in the subsequent G1 via p21 expression. Nat Commun. 2017;8:14728.

47. Perucca P, Cazzalini O, Madine M, Savio M, Laskey RA, Vannini V, et al. Loss of p21 CDKN1A impairs entry to quiescence and activates a DNA damage response in normal fibroblasts induced to quiescence. Cell Cycle. 2009;8(1):105–14.

48. Tumpel S, Rudolph KL. Quiescence: Good and Bad of Stem Cell Aging. Trends Cell Biol. 2019.

49. Yao G. Modelling mammalian cellular quiescence. Interface Focus. 2014;4(3):20130074.

50. Hanahan D, Weinberg Robert A. Hallmarks of Cancer: The Next Generation. Cell. 2011;144(5):646–74.

51. Andreini C, Banci L, Bertini I, Rosato A. Counting the zinc-proteins encoded in the human genome. J Proteome Res. 2006;5(1):196–201.

52. Lambert SA, Jolma A, Campitelli LF, Das PK, Yin Y, Albu M, et al. The Human Transcription Factors. Cell. 2018;172(4):650–65.

53. Danielsen JR, Povlsen LK, Villumsen BH, Streicher W, Nilsson J, Wikstrom M, et al. DNA damage-inducible SUMOylation of HERC2 promotes RNF8 binding via a novel SUMO-binding Zinc finger. J Cell Biol. 2012;197(2):179–87.

54. Ahel I, Ahel D, Matsusaka T, Clark AJ, Pines J, Boulton SJ, et al. Poly(ADP-ribose)-binding zinc finger motifs in DNA repair/checkpoint proteins. Nature. 2008;451(7174):81–5.

55. Witkiewicz-Kucharczyk A, Bal W. Damage of zinc fingers in DNA repair proteins, a novel molecular mechanism in carcinogenesis. Toxicol Lett. 2006;162(1):29–42.

56. Prasad AS, Beck FW, Snell DC, Kucuk O. Zinc in cancer prevention. Nutr Cancer. 2009;61(6):879–87.

57. Pan Z, Choi S, Ouadid-Ahidouch H, Yang J-M, Beattie JH, Korichneva I. Zinc transporters and dysregulated channels in cancers. Front Biosci (Landmark Ed). 2017;22:623–43.

58. Chandler P, Kochupurakkal BS, Alam S, Richardson AL, Soybel DI, Kelleher SL. Subtype-specific accumulation of intracellular zinc pools is associated with the malignant phenotype in breast cancer. Molecular Cancer. 2016;15(1):2.

59. Debnath J, Muthuswamy SK, Brugge JS. Morphogenesis and oncogenesis of MCF-10A mammary epithelial acini grown in three-dimensional basement membrane cultures. Methods. 2003;30(3):256–68.

60. Carter KP, Carpenter MC, Fiedler B. Critical Comparison of FRET-Sensor Functionality in the Cytosol and Endoplasmic Reticulum and Implications for Quantification of Ions. Anal Chem. 2017;89(17):9601–8.

