## Supplementary Note for "Single Cell Analysis Reveals Multiple Requirements for Zinc in the Mammalian Cell Cycle"

### 1. Automated cell segmentation and tracking

To analyze imaging data in a systematic and high throughput way, we developed an automated image analysis pipeline. The basic steps are summarized in Figure 1 and consist of segmentation or identification of individual cells by their nuclei, measurement of cell data, and tracking or associating the data collected in the current frame to individual cells. Details are provided in the following sections. All code was developed in MATLAB (R2018a).

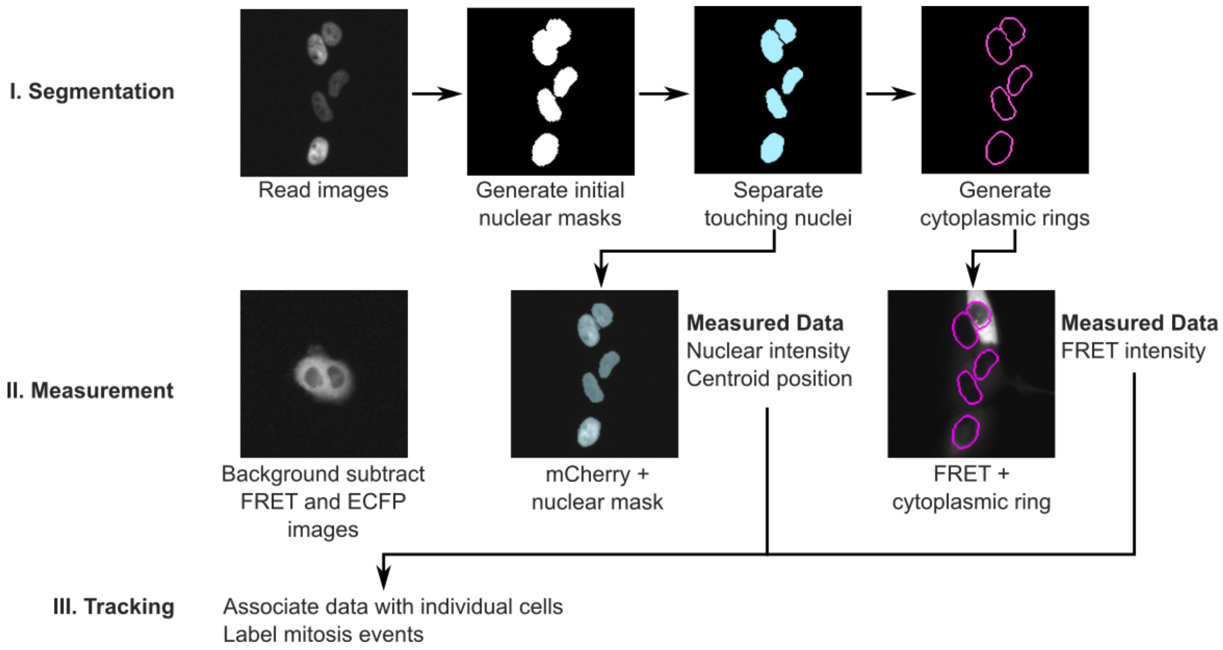


**Figure 1:** Schematic of the automated image analysis pipeline. **(I) Segmentation:** First, the cell nuclei are segmented (identified) using an intensity threshold, obtained from the image intensity histogram. Clumps of nuclei were then separated using the watershed algorithm to obtain the nuclear masks. To measure the cytoplasmic intensity, rings were generated by dilating then subtracting the nuclear masks. **(II) Measurement:** In the measurement step, the fluorescence images (for example FRET and CFP images) were first background-subtracted. The background intensity was estimated by dividing the image into 11x11 blocks. The lowest fifth percentile intensity value for each block was then computed and used as the background. The nuclear intensity was then measured using the nuclear mask, while the intensities in the individual fluorescence channels (e.g. FRET and ECFP channels) were measured using the cytoplasmic rings. The resulting FRET ratio (FRET/CFP) was then computed. The position of each cell was also measured as the centroid of the nuclear masks. **(III) Tracking:** The measured cell data was then assigned to tracks representing individual cells. This process was repeated for each frame in the movies.

### 1.1 Segmentation

#### 1.1.1 Generating the nuclear mask

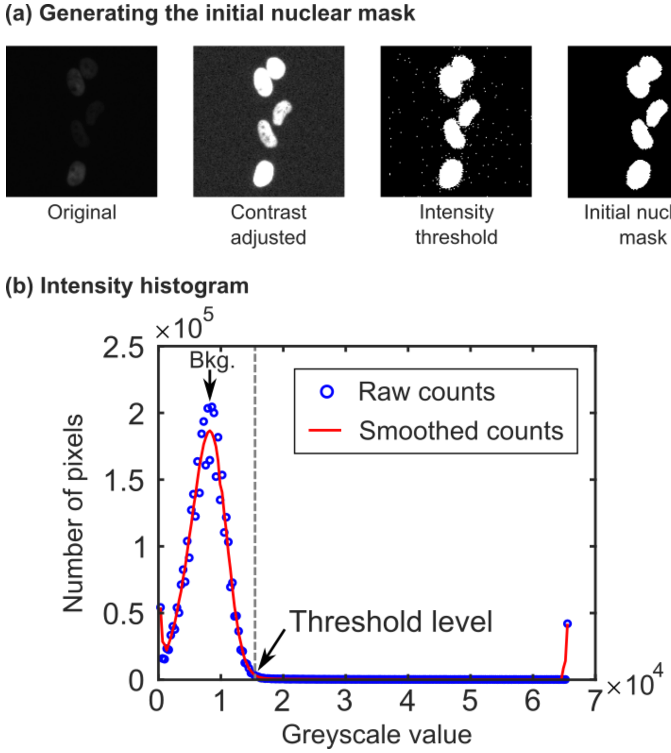


**Figure 2:** (a) Representative images showing segmentation of the nuclei. The contrast of the raw nuclear image was adjusted to better identify the nuclei. An intensity histogram of the contrast-adjusted image was then computed (b). The intensity counts were smoothed, then the first peak was identified as the background (*Bkg.*). The threshold intensity was then chosen to be the intensity where the number of counts dropped to 10% of the background counts. An intensity threshold was then applied to generate a mask. Small spots were removed and the edges of the mask were smoothed using morphological opening.

In our algorithm, individual cells are identified by their nuclei, which are tagged with a fluorescence marker (H2B-mTurquooise, H2B-mCherry or H2B-Halo tag). In the first step, the individual nuclei are segmented (identified) as shown in **Figure 2**. The goal of this step is to generate a binary *mask*, which is a matrix the same size as the image that has a value of 1 for pixels containing cell nuclei and 0 for pixels containing background.

First, the contrast of the raw image data was adjusted to assist with nucleus identification. The pixel values in the raw image were stretched linearly using the equation

$$I_{cs} = \left( \frac{I_{raw} - P_5}{P_{95} - P_5} \right) \tag{1}$$

where $I_{cs}$ is the contrast-stretched pixel intensity, $I_{raw}$ is the raw pixel intensity, and $P_5$ and $P_{95}$ are the greyscale values corresponding to the 5-th and 95-th percentile of intensity values in the raw image. Note that this contrast-stretched image was only used to generate the nuclear mask.  All intensity values reported were measured using the raw images.

An initial nuclear mask was then generated by selecting pixels in the contrast-adjusted image above a threshold value. The threshold intensity was identified by calculating the intensity histogram of the image, shown in **Figure 2**(b). The background intensity distribution was identified as the peak occurring at the lowest intensities (labelled *Bkg.* in the figure). The threshold was then selected to be the intensity where the pixel counts first drops to less than 10% of the maximum counts of the background.

The edges of the nuclear mask were then smoothed by morphological opening with a disk-shaped structuring element with a diameter of 2 pixels. Regions with less than 50 pixels were also removed to remove artifacts caused by image noise.

The initial nuclear mask generated typically contains nuclei which are touching. To separate these nuclei, we used the watershed algorithm (Meyer, 1994). Our process is outlined in **Figure 3**. The watershed algorithm treats an image as a topographical map, with intensity as height. The algorithm then finds the lines between adjacent "basins".

To generate the topographical representation needed for the watershed algorithm, we used a distance transform to label the pixels in the mask with the distance to the edge of each nucleus. These distances are then made negative so the center of each cell would be "deepest". The background pixels were then set to –Inf. The distance transform operation is illustrated in **Figure 3** (b).

To avoid over-segmentation (i.e. a single cell is mislabeled as multiple cells), local minima with a depth of less than 1.2 were suppressed. We then used the built-in MATLAB function "watershed" to determine the border between adjacent nuclei, as shown in **Figure 3** (a). Finally, any nuclear masks intersecting the edge of the image were removed to avoid tracking cells which were only partially in the field-of-view.

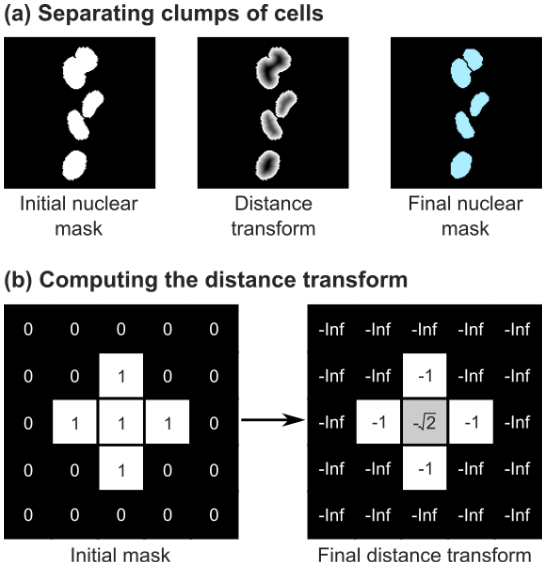


**Figure 3:** Separating clumps of cells. (a) Representative images showing the nuclear mask, distance transform, and final nuclear mask after the watershed algorithm. (b) Schematic showing the distance transform operation.

##### 1.1.2 Generating the ring masks

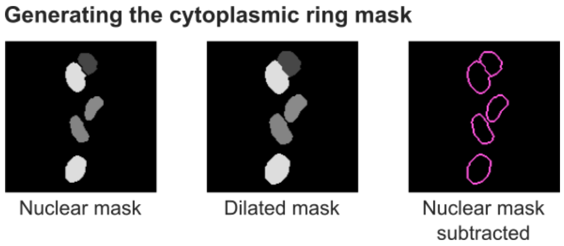


**Figure 4:** Schematic showing the generation of the cytoplasmic rings used to measure the FRET and ECFP intensities.

In general, the cytoplasm of each cell is indistinguishable in the fluorescence channels as the cell membranes were not tagged. Hence, we followed the approach used by (Spencer et al., 2013) and measured cytoplasmic intensity in a ring around the nucleus.

As shown in **Figure 4**, to generate these cytoplasmic rings, the nuclear masks are first dilated by 2 pixels. The nuclear masks are then subtracted, leaving a ring around the nucleus.

#### 1.2 Measurements

For each individual, unconnected object in the nuclear mask, the following properties were measured.

##### 1.2.1 Nuclear intensity and centroid position

The nuclear intensity of a cell is measured as the mean intensity of the H2B channel within its associated nuclear mask. Representative images and plots of the nuclear intensity are shown in **Figure 8** (a). The centroid (or center-of-mass coordinate) of the nucleus mask was also recorded. These coordinates were subsequently used in the cell tracking algorithm to identify individual cell data.

##### 1.2.2 FRET ratio for measuring $Zn^{2+}$ using NES-ZapCV2

The FRET ratio was computed using [need citation]

$$\text{FRET ratio} = \frac{\text{FRET} - \text{FRET}_{bg}}{\text{ECFP} - \text{ECFP}_{bg}} \tag{2}$$

where the subscript *bg* refers to the background intensity value for the FRET and ECFP channels respectively.

To determine the background intensity, we divided the image into 11x11 sub-images. The intensity value corresponding to the lowest 5-th percentile of each sub-image was then computed and used as the background, as shown in **Figure 5**(a). This operation was repeated for the FRET and ECFP channels separately.

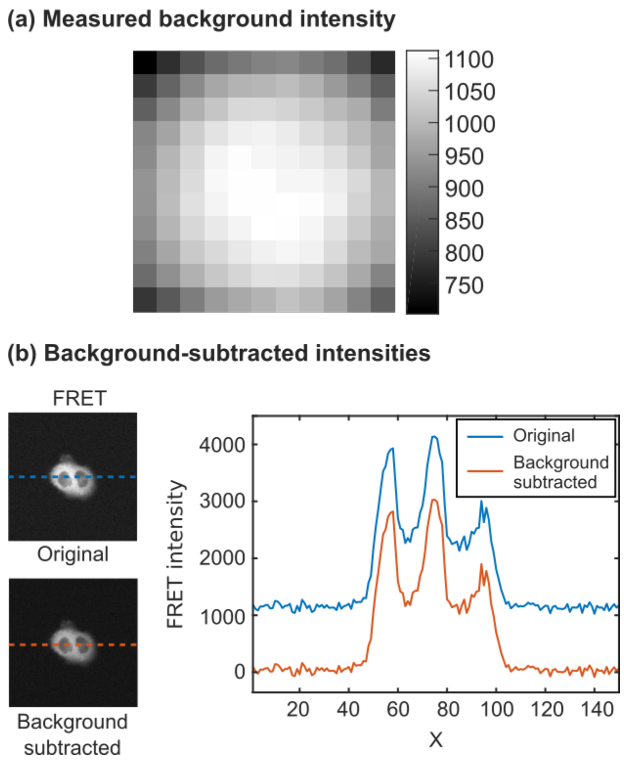


**Figure 5:** Measurement of image background. (a) The computed background intensities for each of the 11x11 sub-images. (b) Representative images of a cell in the FRET channel, showing the original image and the background-subtracted image. The intensity profiles along the dotted lines are shown on the right.

Since the FRET signal here is nuclear-excluded, the FRET and ECFP intensities in the cytoplasm were measured as the mean intensity within the ring mask. The resulting FRET ratio was then computed using Eq. 2. Representative images showing the FRET ratio are shown in **Figure 8**(b).

1.2.3 CDK2 ratio for defining CDK2 activity throughout the cell cycle

In the experiments using the CDK2 sensor, the CDK2 ratio is defined as

$$\text{CDK2 ratio} = \frac{\text{CDK2}_{cyto} - \text{CDK2}_{bg}}{\text{CDK2}_{nucl} - \text{CDK2}_{bg}} \tag{3}$$

where $\text{CDK2}_{\text{cyto}}$ is the mean intensity of the CDK2 channel within the cytoplasmic ring, and $\text{CDK2}_{\text{nucl}}$ is the mean intensity within the nuclear mask. The CDK2 images were background-corrected in the same manner as the FRET and ECFP images as described in 1.2.2 above. Representative images showing the measured CDK2 ratio are shown in **Figure 9**.

### 1.3 Cell tracking algorithm

Tracking refers to the task of collecting data belonging to individual cells. To track individual cells, we used an algorithm based on the linear assignment method (Jaqaman et al., 2008). The linear assignment problem is a classic problem in computer science: say there are a number of workers and tasks. A cost matrix, which contains the cost of assigning a worker to a particular task is provided. The problem is to assign each worker a task, while minimizing the total cost.

Following Jaqaman's framework, we can modify this problem for cell tracking: we now need to assign each cell detected in the current frame with cells detected in the previous frames. However, not every cell will be present in either frame due to cells leaving or entering the field-of-view or cells dividing. Hence, we also have to provide the possible outcomes that cells will not be linked.

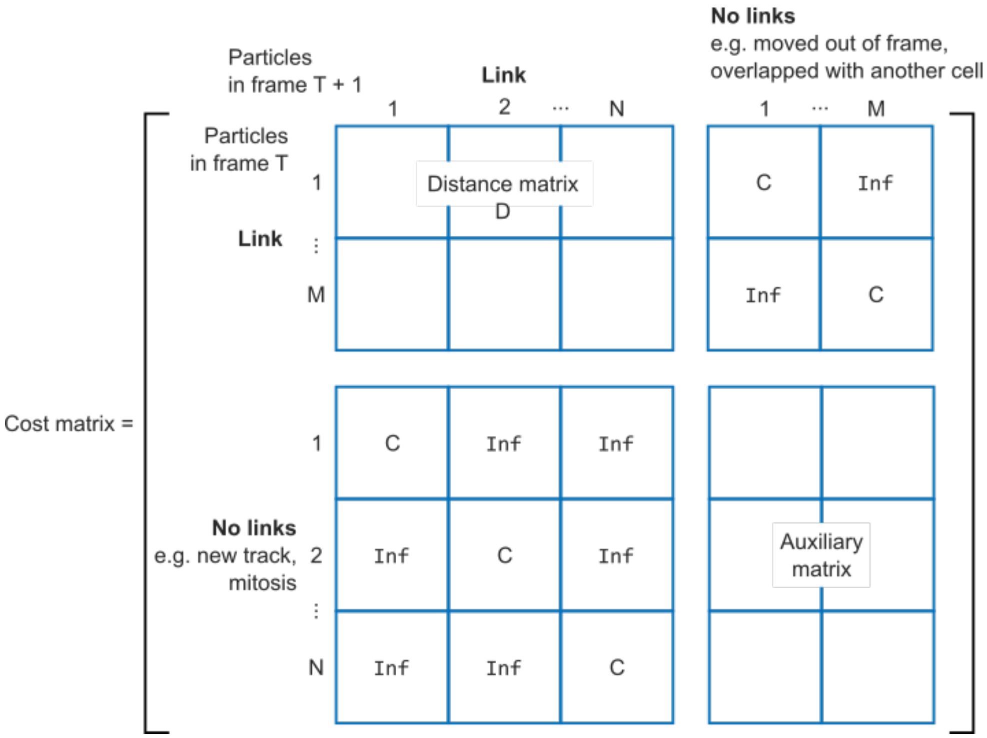


**Figure 6:** Components of the cost matrix used for linking cell data to tracks. The linking algorithm used was based on (Jaqaman et al., 2008). M is the number of existing tracks, representing individual cells in the movie. N is the number of cells detected in the current frame T + 1. The blocking value of infinity (Inf) is assigned to elements in the matrix where assignments must never occur.

Hence, the cost matrix can be broken into four quadrants, as shown in **Figure 6**. The top-left quadrant contains the cost matrix D for linking a cell in the current frame with a cell in the previous frame. This cost is defined as the Euclidean distance between the last position of cells detected in the previous frame to the position of all detected objects in the current frame. To avoid linking cells across physically impossible distances, we set a blocking value of infinity (Inf) for distances greater than 100 pixels. This distance was determined manually from the collected images.

The top-right quadrant contains the cost for cells in the previous frame to have no links with cells in the current frame. Assignment of tracks to these columns indicate that the cell has drifted out of the field-of-view, or has otherwise been lost (overlapping or touching another cell).

Similarly, the bottom-left quadrant represent the cost of cells in the new frame not being assigned to an existing track. If cells are assigned to this outcome, they represent new cells generated either due to a mitosis event or entered the field-of-view along the edges of the image. The value for the alternative cost C is defined as $1.05 \times \max(D)$.

The bottom-right quadrant contains the auxiliary matrix, used only to complete the matrix. This matrix was set to be the transpose of the linking cost D, with all non-blocking costs replaced by the minimum cost in D.

The cost matrix assignments are then solved using the Jonker-Volgenant algorithm (Jonker and Volgenant, 1987). After assignment, each row is assigned to a column.

**Linking.** Linking occurs when a column (representing a cell in the current frame) is assigned to the first M rows.

**Stop tracking or missing frame.** Rows which are assigned to column beyond the first N columns represent cells which are missing in the current frame. Assignment of tracks to these columns indicate that the cell has drifted out of the field-of-view, or has otherwise been lost (overlapping or touching another cell).

**Detecting mitosis events.** To test if a cell had divided, nuclei in a region of 30 pixels around each unassigned nucleus are identified. If there is an existing nucleus with a similar area as the unassigned nucleus (i.e. within 30%), has a similar intensity (within 10%) and that nucleus had not divided recently (within the last 2 frame), then a mitosis event is recorded. In this case, tracking is stopped for the mother cell and two new tracks are created for the daughter cells.

### 1.4 Image registration

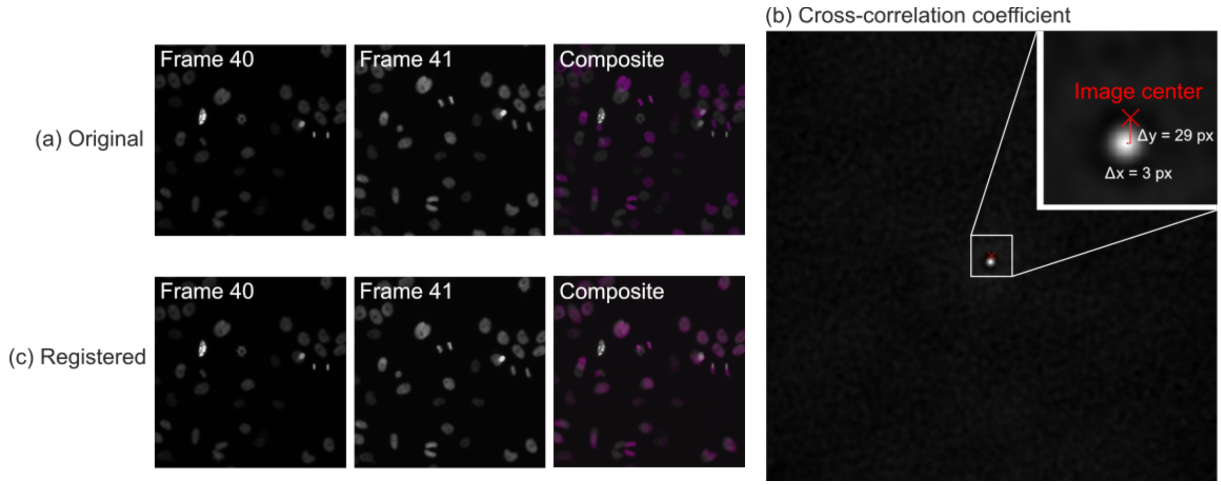


**Figure 7:** Image registration was used to correct for image displacements occurring when zinc was resupplied (between frames 40 and 41). (a) In the original images, the displacement is easily seen in the composite image (frame 40 in greyscale; frame 41 in magenta). Note that some cells have moved and started dividing between the frames. (b) To determine the lateral shift in the image, the cross-correlation coefficient was computed. The shift between the images is the distance from the maximum of the cross-correlation coefficient to the image center. (c) The lateral shift is then applied to frame 41 and beyond.

In the datasets where zinc was resupplied, the removal and subsequent replacement of the plate resulted in a lateral translation being introduced. As shown in **Figure 7** (a), there is both a bulk translation of the image as well as movement of the cells. As the tracking code uses position to identify the cells, these shifts result in the wrong cells being assigned.

To correct for this translation, we computed the pixel shift between the images using the cross-correlation of the reference image (i.e. the last frame before the plate was removed for zinc resupply) with the moved image (i.e. the next recorded frame after the plate was replaced) as:

$$
\begin{aligned}
K &= I_{ref} * I_{moved} \\
&= \mathcal{F}^{-1}\left\{\mathcal{F}\{I_{ref}\} \times \mathcal{F}\{(I_{moved}\}\right\}
\end{aligned}
$$

where F refers to the Fourier transform and * refers to the convolution operation. The shift between the images is then obtained by measuring the distance from the maximum of the cross-correlation coefficient to the image center, as shown in **Figure 7** (b). Frames after zinc resupply were then shifted by this amount, as shown in **Figure 7** (c).

### 2. Representative images

The following Figures **Figure 8** and **Figure 9** show representative images, masks, and tracked intensities from the image analysis and cell tracking code.

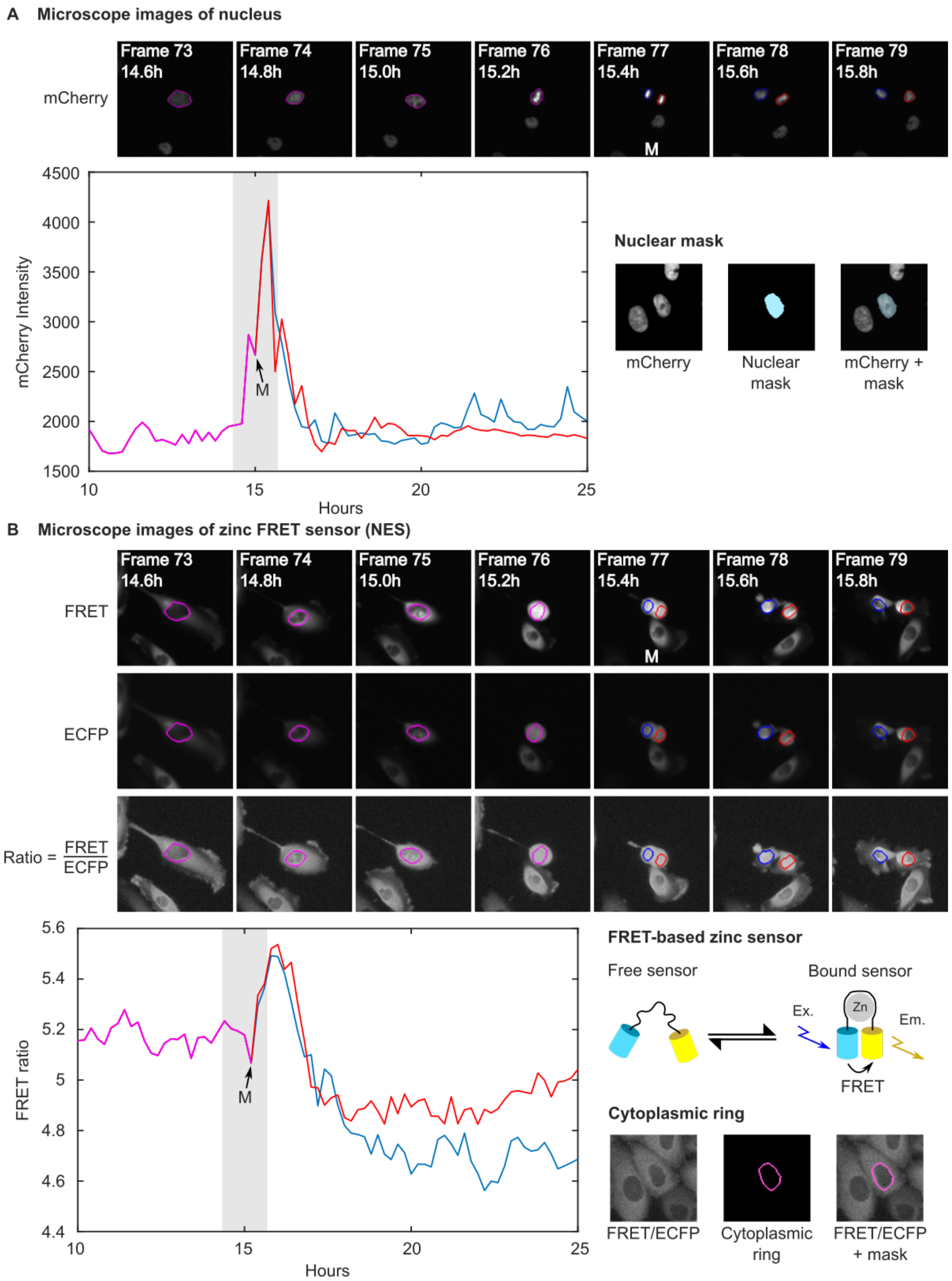


**Figure 8:** (a) shows how the nuclear (H2B-mCherry) intensity was measured. The nuclear masks are shown in the right inset. M indicates the frame at which mitosis occurs (defined as the frame when the nucleus splits into two distinct objects). The shaded region of the intensity plot indicates when the image

stills were taken. (b) The corresponding images acquired in the FRET and ECFP channels, as well as the computed FRET ratio, are shown.

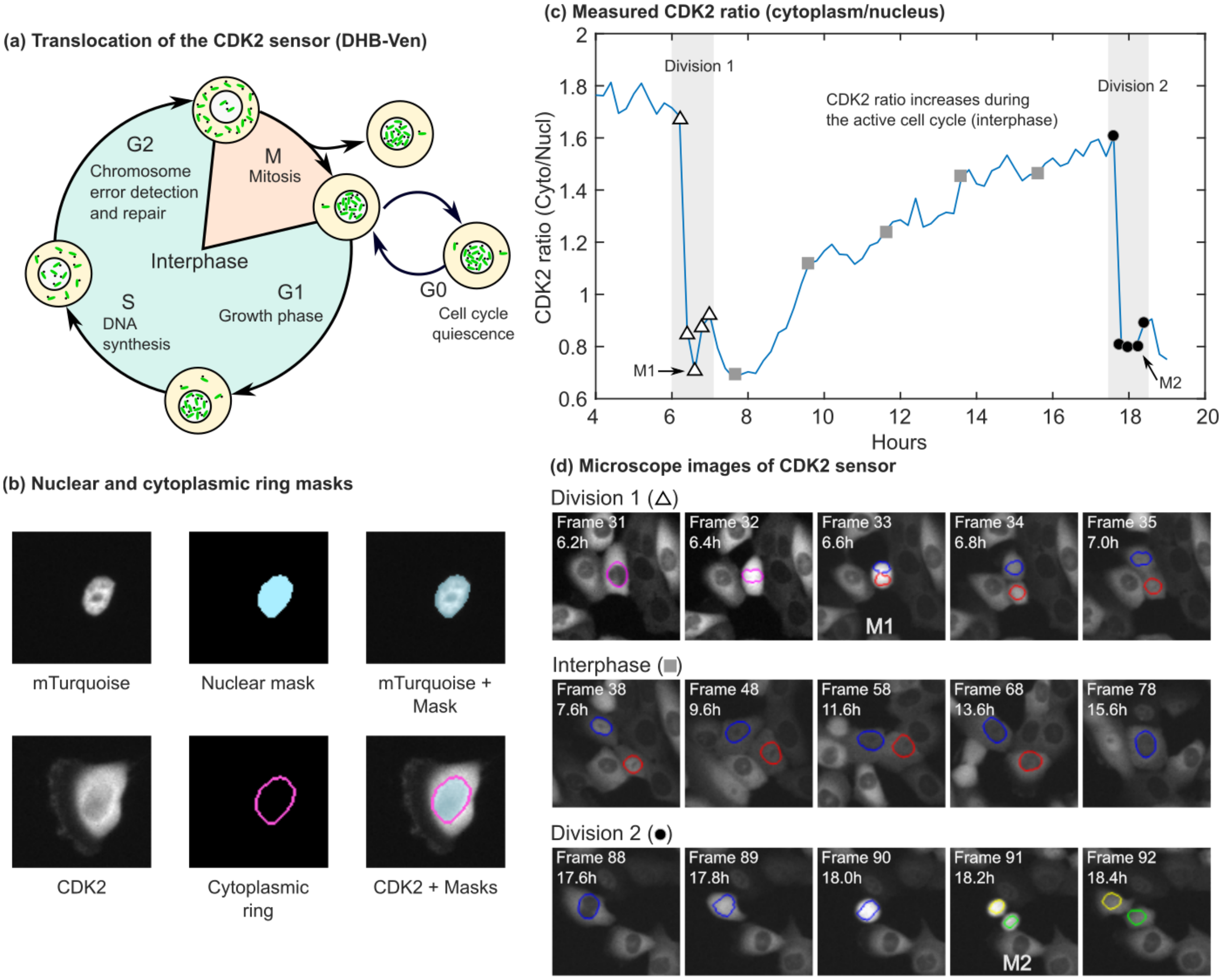


**Figure 9:** (a) Schematic illustrating the location of the CDK2 sensor at different points within the cell cycle (adapted from Spencer, 2013). The CDK2 ratio, defined as the ratio between the intensity within the cytoplasm and the nucleus, is lowest during mitosis M, then builds up during G1 – S and is highest during the G2 phase. (b). The nuclear and cytoplasmic ring masks used to measure intensity of the CDK2 intensity within the nucleus and cytoplasm. Here, H2B-mTurquoise was used for nuclear tracking. (c) Representative trace of the CDK2 ratio showing two cell division events (shaded regions). (d) Microscope images corresponding to the CDK2 ratio plot in (c). The corresponding data points are marked on (c) with the white triangle, the grey square and the black circles.

#### 3. Accuracy of segmentation

To assess the accuracy of cell segmentation, we manually identified errors from three frames of a representative movie (Well A08, CDK2 dataset). This movie was chosen as the cells were proliferative and became crowded at the end of the movie, representing the most challenging condition for segmentation.

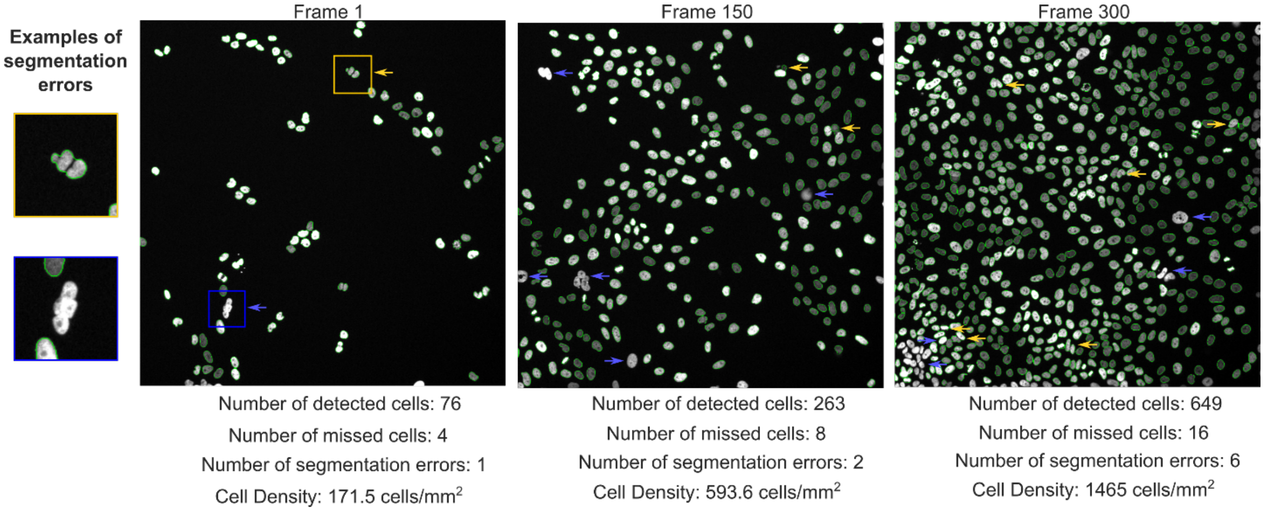


**Figure 10:** Characterizing the errors in segmentation. The outline of each cell identified by the segmentation code is drawn in green. The blue arrows indicate missing cells, while the yellow arrows indicate segmentation errors.

We counted two types of errors: the number of missing cells and segmentation errors (i.e. cases where a single cell was segmented into multiple parts, cases where multiple cells were not separated, or cases where the watershed algorithm split cells inaccurately). As can be seen in **Figure 10**, the total number of errors increases as the cell density increases.

If we define an error rate as the ratio of the sum of the number of missed cells and the number of segmentation errors over the total number of detected cells, the error rate due to segmentation is between 4 – 7%.

A movie showing the identified mitosis events is included as **SI Movie 1**. In the video, mitosis events identified by the code are indicated by a red cell outline.

**REFERENCES**


Jaqaman, K., Loerke, D., Mettlen, M., Kuwata, H., Grinstein, S., Schmid, S.L., and Danuser, G. (2008). Robust single-particle tracking in live-cell time-lapse sequences. Nat. Methods *5*, 695.

Jonker, R., and Volgenant, A. (1987). A shortest augmenting path algorithm for dense and sparse linear assignment problemsEin Algorithmus mit kürzesten alternierenden Wegen für dichte und dünne Zuordnungsprobleme. Computing *38*, 325–340.

Meyer, F. (1994). Topographic distance and watershed lines. Signal Process. *38*, 113–125.

Spencer, S.L., Cappell, S.D., Tsai, F.-C., Overton, K.W., Wang, C.L., and Meyer, T. (2013). The Proliferation-Quiescence Decision Is Controlled by a Bifurcation in CDK2 Activity at Mitotic Exit. Cell *155*, 369–383.
