## Supplementary Information for "Single Cell Analysis Reveals Multiple Requirements for Zinc in the Mammalian Cell Cycle"

Supplemental Figure 1 …………………………………………………………..2

Supplemental Figure 2 …………………………………………………………..3

Supplemental Figure 3 …………………………………………………………..4

Supplemental Figure 4 …………………………………………………………..5

Supplemental Figure 5 …………………………………………………………..6

Supplemental Figure 6 …………………………………………………………..7

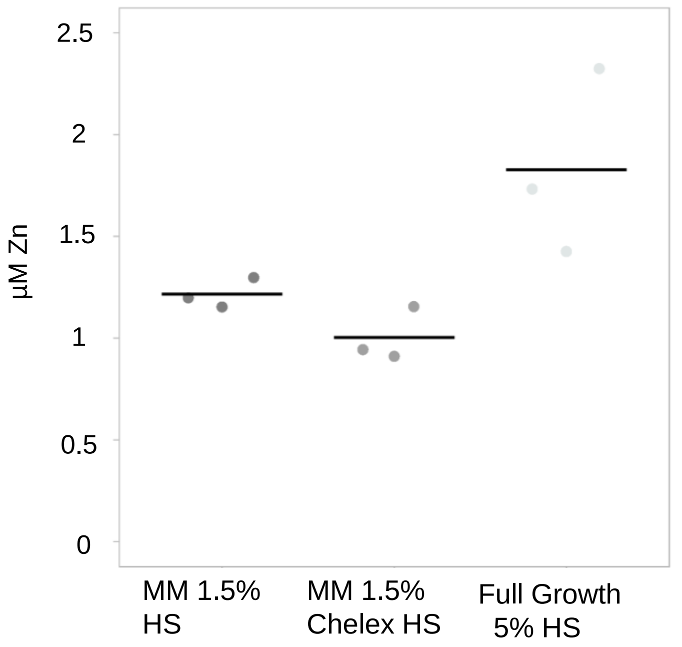


**Supplemental Figure 1. Total Zn content in growth media**
ICP-MS analysis was used to measure Zn content in Minimal Media containing 1.5% horse serum. MM was further treated with Chelex to remove additional Zn. Full growth media containing 5% horse serum was included for reference. n=3 replicates per condition and black bars indicate mean.

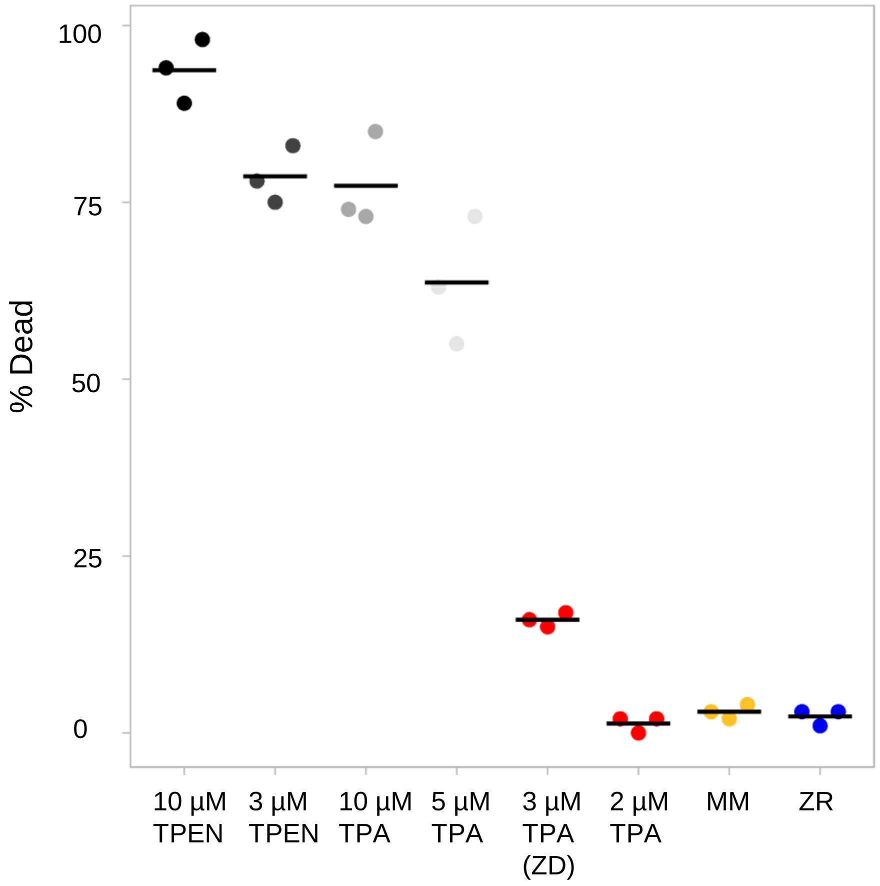


**Supplemental Figure 2. TPEN induces high levels of cell death, while TPA at low levels causes significantly less cell death.**
% dead cells after 30 hours of growth in defined medias. Cell death was measured using trypan blue and cells were counted using an automated cell counter. n=3 per condition. Black bars indicate mean for each condition.

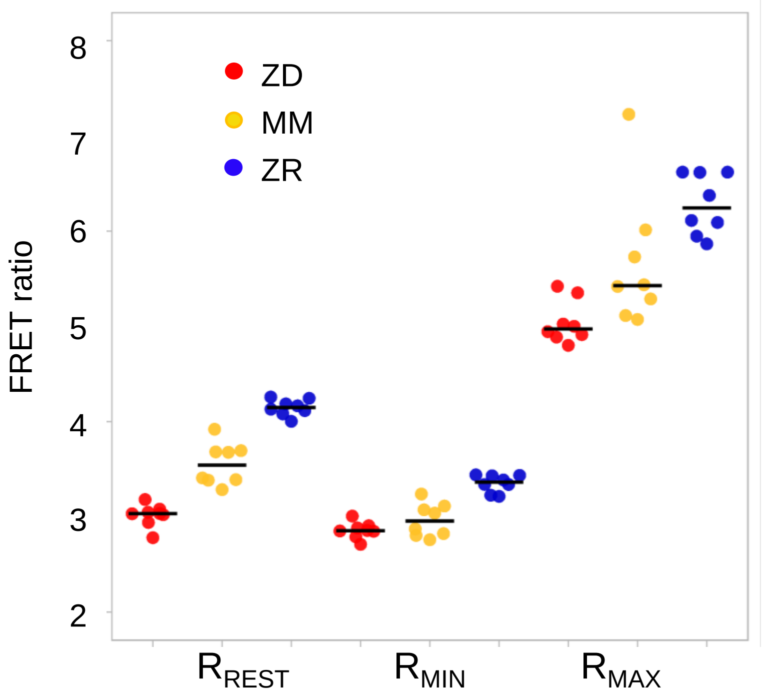


| | ZD | MM | ZR |
|---|---|---|---|
| FS, % | 7.3 ± 0.6 | 22.3 ± 1.0 | 27.4 ± 1.4 |
| DR | 1.77 ± 0.03 | 1.9 ± 0.05 | 1.87 ±0.02 |
| [$Zn^{2+}$], pM | 1 ± 0.2 | 81 ± 29 | 213 ± 51 |

**Supplemental Figure 3. Intracellular $Zn^{2+}$ levels titrate with extracellular media.** Sensor calibrations of MCF10A cells stabling expressing PB-NES-ZapCV2. Cells were grown for 24 hours in either ZD (3µM TPA), MM, or ZR media. For collection of Rrest, cells were imaged in ZD, MM, or ZR. To collect Rmin, 50µM TPA in MM was added. Rmax was measured by treatment with HBSS buffer with 119nM $Zn^{2+}$, 0.001% saponin, and 5µM pyrithione. Fractional saturation (FS) and dynamic range (DR) of the sensor were calculated. $Zn^{2+}$ concentrations were measured as described in the methods. n=8 per condition and black bars indicate mean.

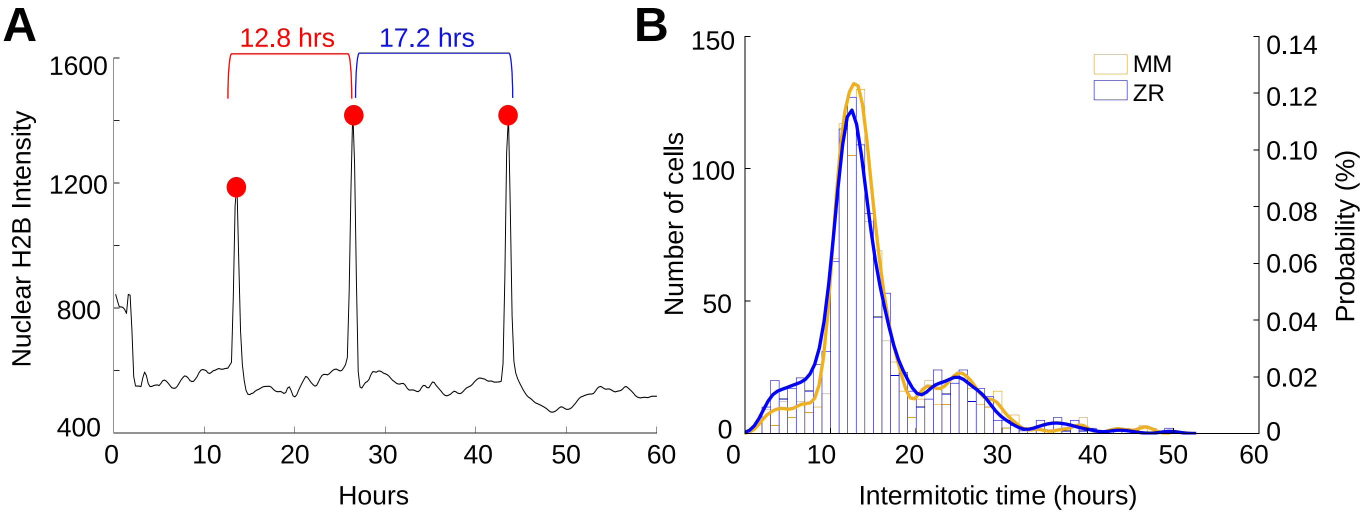


**Supplemental Figure 4. Intermitotic time does not differ between cells grown in minimal media and $Zn^{2+}$ replete conditions.**

**A.** H2B intensity of a single cell tracked throughout a 60 hour period. Three mitosis events were detected and marked with a red dot. The intermitotic time (IM) was calculated by determining the difference in hours between each successive mitosis event and indicated above.

**B.** Intermitotic times for divisions within a 60 hour period of cells grown in either MM or ZR conditions. Histograms show number of cells in each IM time bin and overlaid density plots show the probability of IM times, peaking ~13 hours for both conditions (n=907 in MM and n=1000 in ZR).

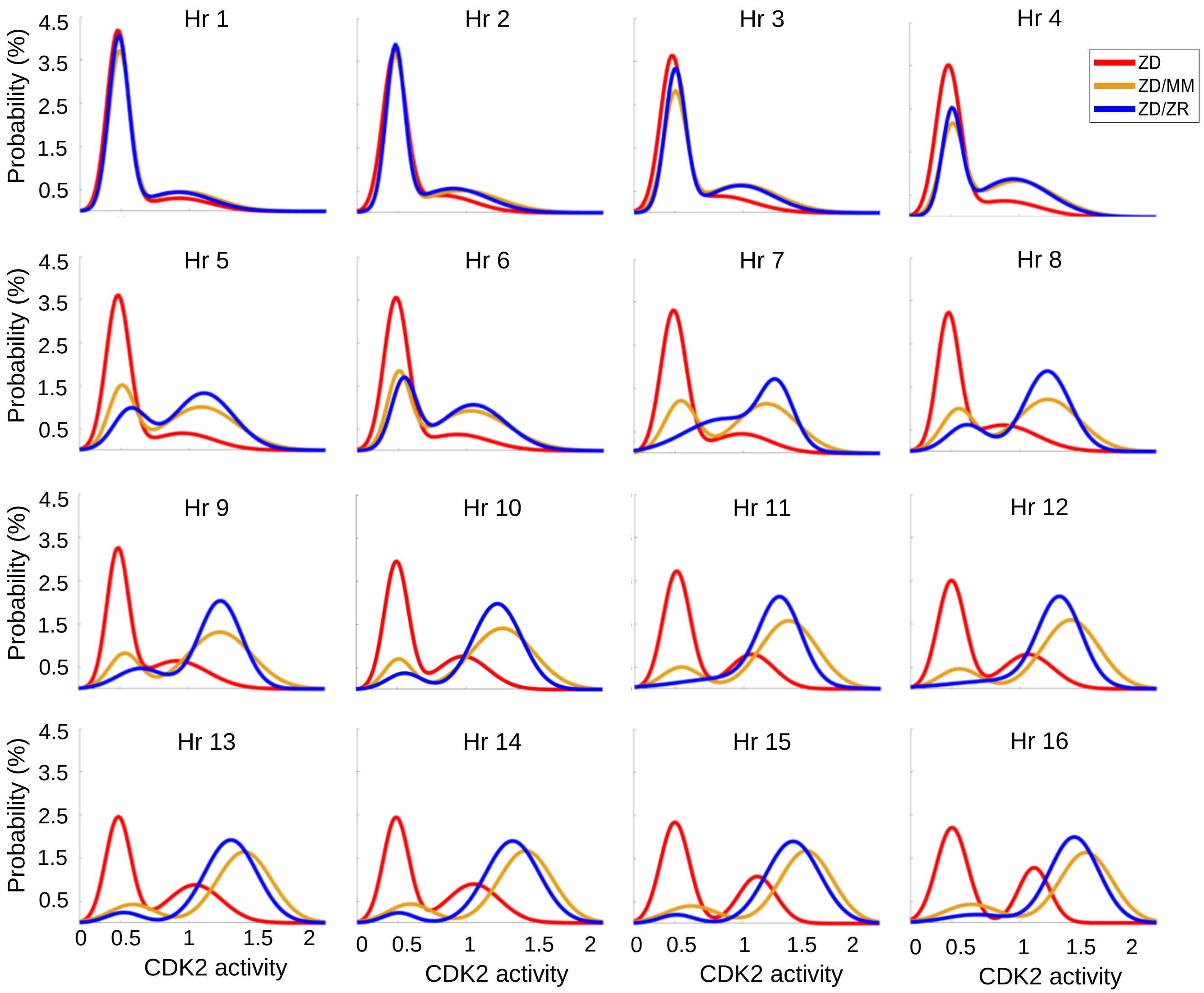


**Supplemental Figure 5. $Zn^{2+}$ resupply allows for re-entry into the cell cycle following $Zn^{2+}$ deficiency-induced quiescence.**
CDK2 activity density plots after cells either remain in ZD media or are resupplied with MM or ZR. Hr indicates number of hours after resupply, e.g. Hr 1 = 1 hour after resupply, or hour 25 of the experiment (n = 210, 207, 202, 195, 196, 196, 191, 187, 185, 188, 188, 187, 187, 181, 178, and 176, for Hrs 1-16 for ZD pooled from 4 wells, 1068, 1121, 1129, 1124, 1127, 1142, 1149, 1154, 1165, 1169, 1170, 1178, 1191, 1187, 1182, and 1198, for MM pooled from 8 wells, and 1024, 1067, 1085, 1103, 1120, 1124, 1137, 1139, 1144, 1150, 1148, 1153, 1156, 1157, 1166, and 1162 for ZR pooled from 8 wells).

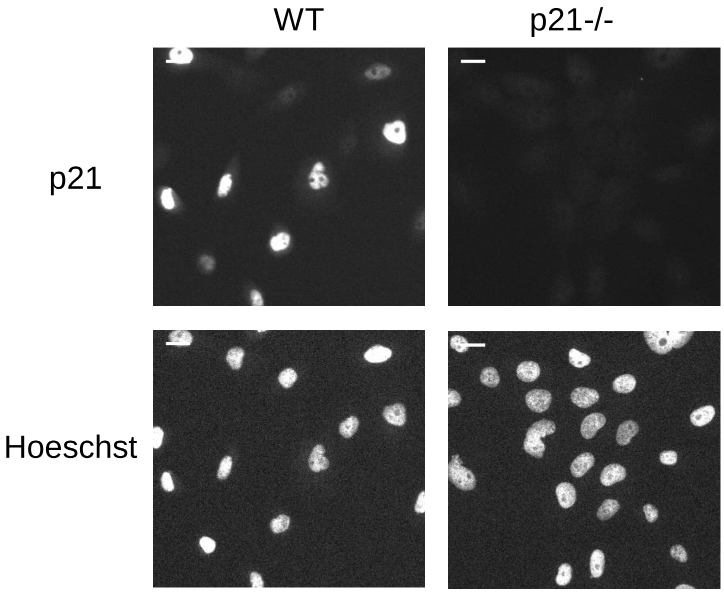


**Supplemental Figure 6. Validation of p21 knockdown in p21-/- cell line.**
Representative cell images for Hoescht DNA staining and p21 IF for WT MCF10A cells and p21-/- MCF10A cells. p21 was detected with an anti-p21 antibody. Scale bars indicate 20 µm.
